# Release, rescue and recycling: termination of translation in mammalian mitochondria

**DOI:** 10.1101/2021.03.10.434893

**Authors:** Eva Kummer, Katharina Noel Schubert, Tanja Schönhut, Alain Scaiola, Nenad Ban

## Abstract

The mitochondrial translation system originates from a bacterial ancestor but has substantially diverged in the course of evolution. Here, we use single particle cryo-EM as a screening tool to identify mitochondrial translation termination mechanisms and to describe them in molecular detail. We show how mitochondria release factor 1a releases the nascent chain from the ribosome when it encounters the canonical stop codons UAA and UAG. Furthermore, we define how the peptidyl-tRNA hydrolase ICT1 acts as a rescue factor on mitoribosomes that have stalled on truncated messages to recover them for protein synthesis. Finally, we present near-atomic models detailing the process of mitochondrial ribosome recycling, to explain how a dedicated elongation factor, mtEFG2, has specialized for cooperation with the mitochondrial ribosome recycling factor to dissociate the mitoribosomal subunits at the end of the translation process. (134 words)

## INTRODUCTION

Mitochondria are eukaryotic organelles of proteobacterial origin. During evolution, they have maintained a small genome that encodes 13 membrane proteins in humans, which form essential parts of the respiratory chain complexes in the inner mitochondrial membrane (IMM). Efficient expression of these few mitochondrial genes is consequently crucial to supply the cell with sufficient energy and is realized by a specialized mitochondrial translation system. The mitochondrial translation machinery is very different from its cytosolic or bacterial counterparts and shows an enormous variability between eukaryotes with respect to the composition and architecture of the mitochondrial ribosome.(Greber and Ban, 2016) Polypeptide synthesis on the ribosome is coordinated by a number of translation factors that interact with the mitoribosome, the tRNAs and the mRNA during all stages of the translation cycle. Similar to the mitochondrial ribosome itself, the set of translation factors in mitochondria are strikingly diverse in comparison to their well-studied bacterial counterparts.(Ott et al., 2016)

When the ribosome reaches the end of an open reading frame on a mRNA, translation is terminated by positioning of a stop codon into the ribosomal A site. This codon is recognized by ribosomal release factors that couple stop-codon recognition with hydrolysis of the ester bond between nascent chain and P site tRNA, thereby releasing the nascent polypeptide from the ribosome. Bacteria possess two release factors with overlapping specificities, RF1 recognizes UAA and UAG stop codons while RF2 is able to decode UAA and UGA, respectively.(Ito et al., 2000; Korostelev et al., 2008; Laurberg et al., 2008; Petry et al., 2005; Weixlbaumer et al., 2008) Mitochondria show alterations to the otherwise universally conserved genetic code with substantial effects on translation termination. In humans, UGA encodes for tryptophan instead of a stop codon.(Barrell et al., 1979) In addition, the open reading frames for mitochondrial CO1 and ND6 mRNAs end with the alternative stop codons AGA and AGG, respectively, whereas these codons specify arginine incorporation in the cytosol.(Anderson et al., 1981) The situation is made even more complex by the fact that mRNAs that encounter the ribosome may occassionally be missing a stop codon due to nucleolytic cleavage, misprocessing of the RNA or premature abortion of transcription. Those mRNAs cause stalling of the ribosome at their 3’ end. In bacteria, multiple elaborate systems rescue the ribosome stalling: trans-translation by tmRNA and SmpB, the alternative release factor ArfB, as well as the alternative release factor ArfA in conjuction with RF2.(Keiler, 2015) These systems recognize the stalled ribosome via a partially empty mRNA channel and subsequently trigger release of the nascent chain in a codon-independent manner. It is conceivable that the mitochondrial translation apparatus must have evolved strategies to rescue stalled ribosomes in the absence of a stop codon. Mitochondrial translation termination would thus be required for: 1. polypeptide release upon recognition of the conventional stop codons UAA and UAG, 2. polypeptide release triggered by the alternative stop codons AGA and AGG, and 3. rescue of stalled ribosomes from mRNAs missing stop codons (Fig. 1A).

**Fig 1.**
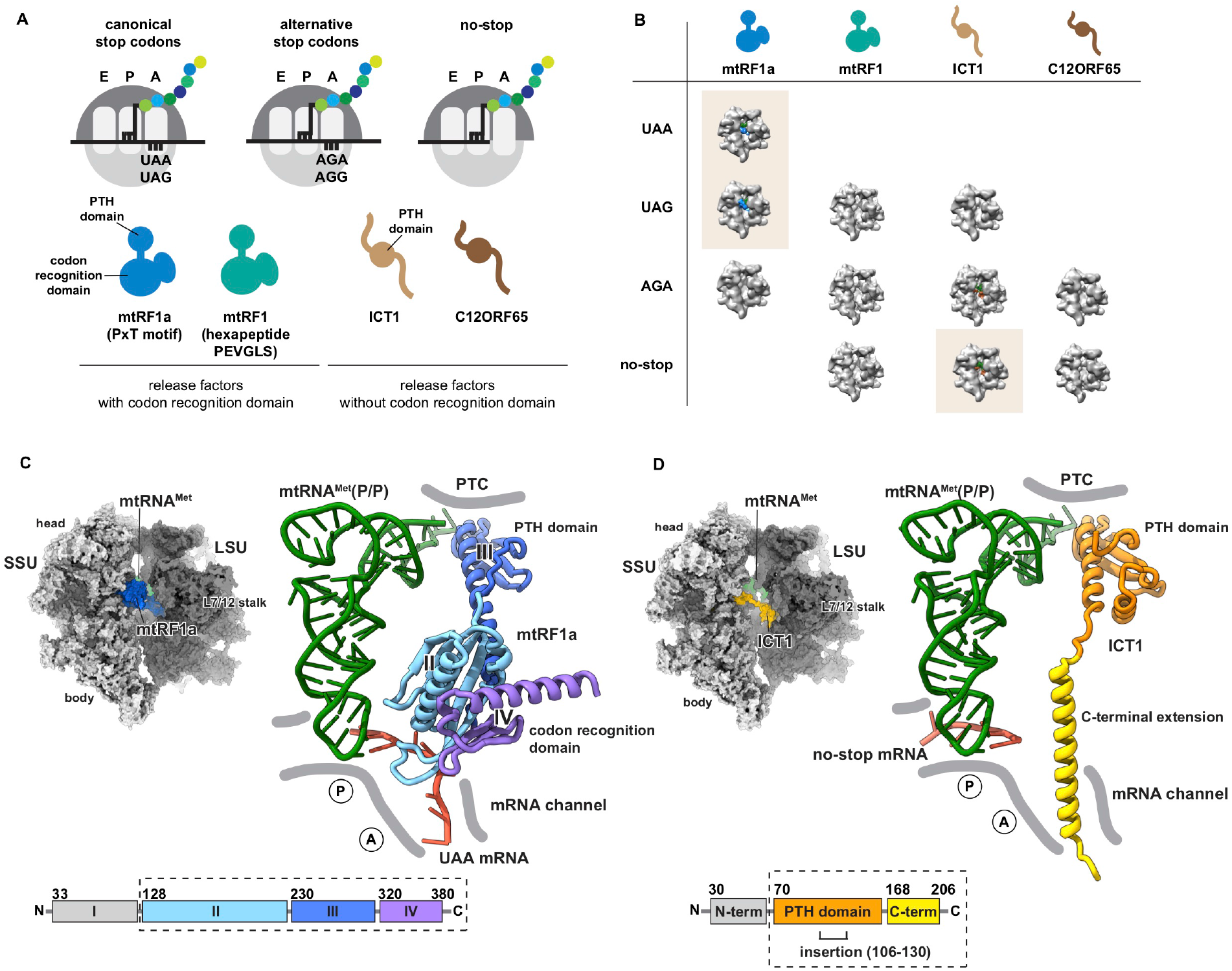
Exhaustive EM screening of translation termination scenarios. A) A schematic overview of the three possible termination scenarios on the mitoribosome are depicted on the top. The ribosomal subunits are shown in grey, mRNA in black with the P site tRNA (black) carrying a polypeptide chain. The four putative translation termination factors in mitochondria are shown below. B) Tabular overview of the tested translation termination conditions. EM densities of the mitoribosome are shown for conditions that have been probed. The empty ribosome is generally shown in grey and binding of a translation termination factor is indicated by colouring of its surface in the EM density (blue in case of mtRF1a and orange in case of ICT1). Complexes that contain termination factors are for clarity highlighted with orange background. C) Overview of the mtRF1a translation termination complex. The structural model of the entire complex is shown as surface representation. Moreover, the ternary complex of mRNA (red), tRNA^Met^ (green) and mtRF1a is shown in isolation and adjacent ribosomal features are indicated. mtRF1a protein domains are indicated by different colours and the corresponding schematic representation of the mtRF1a domain organization is shown below, with numbers indicating residues delineating domain borders. D) Similarly for mtRF1a, the structural model of the translation termination complex containing ICT1 is depicted on the left and the ternary complex of ICT1, mtRNA^Met^ (green) and mRNA (red) is shown in isolation on the right. The ICT1 domain organization is given below.

Four putative translation termination factors have been described in mitochondria based on their homology to known bacterial release factors and the conservation of the essential catalytic GGQ motif. They can be clustered in two classes (Fig. 1A). Class 1 contains mtRF1 and mtRF1a that possess the catalytic peptidyl hydrolase (PTH) domain, a codon recognition domain and an N-terminal domain of unknown function. mtRF1 and mtRF1a differ mostly in the sequence motifs predicted to play a role in specific discrimination of stop codons as well as in length and sequence of their N-terminal domains. Class 2 includes ICT1 and C12ORF65, both of which carry a positively charged C-terminal extension instead of a codon recognition domain while the PTH domain is preserved. They differ by a 25 amino acid insertion only found in the PTH domain of ICT1 that is essential for peptidyl-tRNA hydrolase activity on the ribosome but not for ribosome binding.(Handa et al., 2010; Kogure et al., 2012) Previously, peptidyl hydrolase activity on the 55S monosome has been demonstrated biochemically only for mtRF1a and ICT1.(Akabane et al., 2014; Feaga et al., 2016; Kogure et al., 2014; Nozaki et al., 2008; Richter et al., 2010; Soleimanpour-Lichaei et al., 2007; Wesolowska et al., 2014) While the majority of published data, converges on the recognition of the conventional stop codons UAA and UAG by mtRF1a, there have been diverse interpretations of the function of ICT1.(Akabane et al., 2014; Feaga et al., 2016; Lind et al., 2013; Richter et al., 2010; Wesolowska et al., 2014) Moreover, how the mitochondrial ribosome terminates the alternative stop codons AGA and AGG is still a matter of debate.(Akabane et al., 2014; Feaga et al., 2016; Huynen et al., 2012; Lind et al., 2013; Richter et al., 2010; Temperley et al., 2010; Young et al., 2010) These diverse and in parts contradictory data hamper an understanding of the mechanisms governing mitochondrial translation termination in humans.

Once the polypeptide chain has been released, the mitoribosome needs to undergo a recycling step, in which the large and small ribosomal subunits are split to allow dissociation of the mRNA and remaining tRNAs. Subsequently, the mitoribosome can engage in another round of translation. In many bacterial species the canonical elongation factor (EFG) promotes recycling of the ribosome and in addition catalyzes the movement of the mRNA-tRNA module during translation elongation. In mitochondria these two tasks have been allocated to two distinct proteins. While mitochondrial elongation factor 1 (mtEFG1) catalyzes translation elongation, mtEFG2 cooperates with the mitochondrial recycling factor (mtRRF) to trigger splitting of the mitoribosomal subunits(Rorbach et al., 2008; Tsuboi et al., 2009). Curiously, also a number of bacterial species contain two distinct elongation factors. Although their functional interplay is in most cases not yet conclusively defined, at least in *B. burgdorferi* a similar task distribution as for mitochondrial mtEFG1 and mtEFG2 has been described(Suematsu et al., 2010). Currently, it remains elusive how the two proteins have specialized for their tasks and how interference of one factor with the activity of the other is prevented to secure translation efficiency and mitochondrial function.

Here, we have set out to decipher the molecular basis of human mitochondrial translation termination and recycling using an *in vitro* reconstitution approach coupled with structural studies. Our study includes an exhaustive screen of possible translation termination scenarios on the mitoribosome against all four putative release factors. In accordance with biochemical data, we find mtRF1a engaging the canonical stop codon UAA and UAG whereas ICT1 specifically binds ribosomes stalled on mRNAs lacking a stop codon. To visualize the molecular details of their mode of action, we present near-atomic cryo-EM structures of both translation termination factors bound to the mitoribosome. In addition, we show how mtRRF and mtEFG2 cooperate to split the ribosomal subunits after polypeptide release and establish how mtEFG2 has specialized for the process of ribosome recycling.

## RESULTS

### Extensive screening of possible translation termination scenarios

Biochemical and computational data on the role of different mitochondrial translation termination factors are controversial and hamper the understanding of mitochondrial translation termination. In order to resolve these discrepancies and disentangle the function of mitochondrial translation termination factors, we opted to employ an exhaustive biochemical and structural screening approach. For this, we made use of our previously established *in vitro* reconstitution system to assemble mitochondrial translation complexes that mimic possible translation termination scenarios (Fig. 1A).(Kummer et al., 2018) These complexes were subsequently presented to all four putative mitochondrial translation termination factors. We reasoned that authentic termination factor activity should be accompanied with specific binding of the adequate factor to the ribosome. Screening of the reconstituted complexes using single particle cryo-EM revealed under which conditions factor binding could be observed (Fig. 1B).

Our screen shows that the canonical stop codons (UAA or UAG) are exclusively bound by mtRF1a (Fig. 1B). In agreement with published biochemical data, mtRF1a is however not able to decode the alternative stop codon AGA indicating that the factor is solely dedicated to the recognition of canonical stop codons in mitochondria.(Nozaki et al., 2008; Soleimanpour-Lichaei et al., 2007) The second release factor mtRF1 has an extended codon recognition loop in comparison to mtRF1a. Homology modelling, free energy calculations and computer simulations indicated that it may be involved either in termination at canonical stop codons (UAA and UAG), at alternative stop codons (AGG and AGA) or on mRNAs lacking a stop codon.(Huynen et al., 2012; Lind et al., 2013; Young et al., 2010) However, we do not find mtRF1 bound to the mitochondrial ribosome in any of the possible translation termination scenarios (Fig. 1B). Consistently, none of the biochemical studies detected PTH activity of mtRF1 on the ribosome so far.(Nozaki et al., 2008; Soleimanpour-Lichaei et al., 2007) This may either imply that mtRF1 does not exert a function as translation termination factor or that it may require unidentified cofactors for its action, which we cannot account for in our *in vitro* system.

Previously, ICT1 was reported to exert codon-unspecific peptidyl-hydrolase activity, acting on canonical stop codons, non-canonical stop codons and on ribosomes that lack a codon in the A site.(Richter et al., 2010) ICT1 binds in our reconstituted system specifically ribosomes with an empty A site that resemble a no-stop scenario, in which the ribosome is stalled at the 3’ end of a truncated mRNA. We do not find ICT1 engaged with ribosomes that carry mRNAs with canonical stop codons but have detected some EM density accounting for ICT1 in the sample of mitoribosomes that have been programmed with an mRNA containing the alternative stop codon AGA (Fig. 1B). Although such a binding would be in line with the earlier suggestion that ICT1 acts on non-canonical stop codons, a closer analysis of our cryo-EM dataset revealed that ICT1 was predominantly found in the subpopulation of particles that did not appear to harbor mRNA, whereas mRNA-bound ribosomes were devoid of ICT1. Our observations therefore argue against a role of ICT1 in the recognition of non-canonical stop codons. Finally, we also tested binding of C12ORF65 to our reconstituted translation termination scenarios. C12ORF65 mutations have been correlated in a number of studies with defects in mitochondrial translation leading to assembly defects of the respiratory chain and severe disease phenotypes in humans.(Antonicka et al., 2010; Shimazaki et al., 2012; Spiegel et al., 2014; Tucci et al., 2014; Wesolowska et al., 2015) As C12ORF65 lacks a classical codon recognition domain we restricted our test settings to no-stop scenarios and a possible role in termination of non-canonical stop codons. We do not find C12ORF65 engaged with the mitoribosome under these conditions in line with a recent report that identified C12ORF65 (now termed mtRF-R) to team up with the mitochondrial RNA-binding protein C6ORF203 (MTRES1) in a novel translation rescue pathway under conditions when aminoacylated tRNAs become limiting.(Desai et al., 2020) In this pathway, C12ORF65 acts exclusively on the LSU instead of the 55S monosome. Collectively, this observation and our screening data argue that – despite a similar domain architecture - ICT1 and C12ORF65 rescue stalled mitoribosomes by fundamentally different mechanisms.

### mtRF1a recognizes the canonical stop codon UAA via conserved sequence motifs

To understand the molecular basis for translation termination by mtRF1a and ICT1, we solved their near-atomic structures in the context of the mitochondrial ribosome using single particle cryo-EM (Fig. 1C and 1D). We visualized mtRF1a bound to the mitochondrial ribosome in the presence of mRNA carrying a UAA stop codon in the A site and mtRNA^Met^ in the P site at 3.5 Å (Fig. 1C). The architecture of mtRF1a and its positioning in the ribosomal A site are very similar to bacterial RF1.(Laurberg et al., 2008; Petry et al., 2005) mtRF1a consists of four domains, of which domain II and IV form the codon recognition domain that contacts the mRNA and the ribosomal decoding center on the small ribosomal subunit (SSU) (Fig. 1C). Domain III corresponds to the peptidyl hydrolase (PTH) domain containing the catalytic GGQ motif that is placed adjacent to the CCA end of the P site tRNA_Met_ in the peptidyl transferase center (PTC) of the large ribosomal subunit (LSU). In analogy to bacterial RF1, the N-terminal domain I is rather flexible and does not contact the ribosome.(Korostelev et al., 2008; Petry et al., 2005) Due to its low resolution, domain I could not be fitted unambiguously into its corresponding density and was omitted from the model.

UAA stop codon recognition by mtRF1a involves a mixture of steric restrictions, stacking interactions and hydrogen bonding interactions of mtRF1a domain II and the 12S rRNA with the mRNA via elements that are also conserved in bacteria (Fig. 2A and 2B, Fig. S1A). The mtRF1a codon recognition loop that contains the PxT motif cooperates with the tip of α-helix 5 (α5) of the mtRF1a domain II to decipher bases 1 (U) and 2 (A) of the stop codon. α5 restricts the space for the first position nucleotide to a pyrimidine base and hydrogen bonding with conserved Glu141 and Gly138 favors uridine over cytidine in the first position of the codon.(Laurberg et al., 2008) The second position adenine is stacked between the uridine at position 1 and His215 of mtRF1a and hydrogen bonds with Thr208 of the ^206^PxT^208^ motif. The third nucleotide is splayed out to stack on top of G256 (G530 in bacteria) where it engages with Thr216 and Glu203 of mtRF1a. These recognition elements are entirely preserved from the bacterial system highlighting their essential roles for translation termination fidelity (Fig. S1A). Strikingly, this indicates that despite a substantial evolutionary diversification of the mitochondrial translation system recognition of canonical stop codons involves crucial interactions, none of which could be easily supplemented by alternative approaches during evolution.

**Fig 2.**
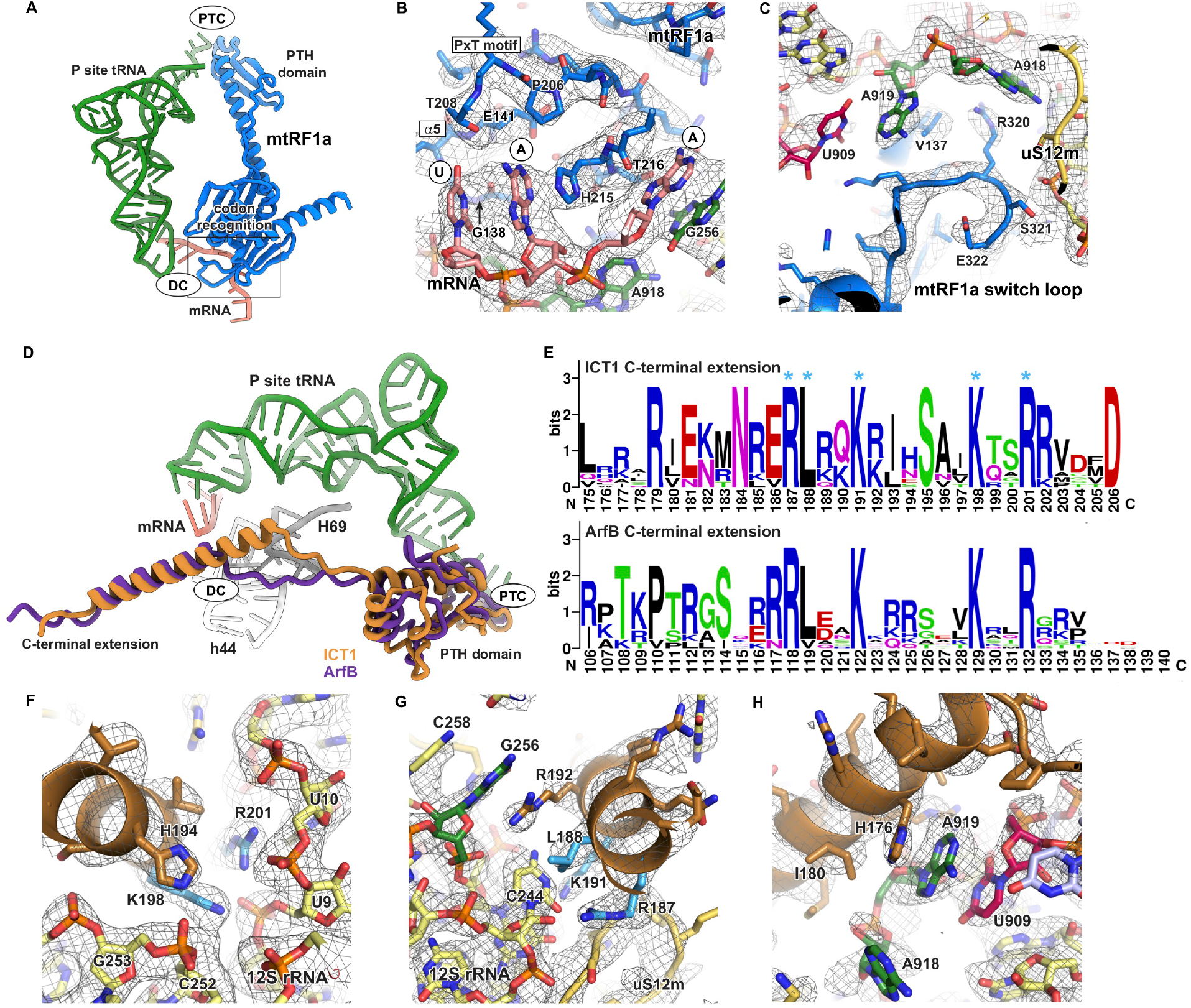
mtRF1a and ICT1 recognize ribosomal complexes via distinct features. A) Overview of the ternary complex of mtRF1a (blue), tRNA (green) and mRNA (salmon) in isolation. Adjacent ribosomal sites are indicated and the interaction site of mtRF1a with the mRNA stop codon is highlighted with a box. PTC = peptidyl-transferase center, DC = decoding center B) mtRF1a (blue) recognizes the UAA stop codon (salmon) via conserved sequence motifs including its _206_PxT_208_ motif and the tip of α-helix 5 (α5). In addition, the second and third base of the stop codon engage in stacking interaction with H215 of mtRF1a and the decoding nucleotide G256 (green) of the 12S rRNA, respectively. The corresponding EM density is shown as mesh. C) The switch loop of mtRF1a is embedded in between the decoding nucleotides of the small ribosomal subunit (green) and the large ribosomal subunit rRNA (red) as well as uS12m (yellow). In contrast to bacteria, the tip of helix 69 (U909, purple) is substantially shorter in mitochondrial ribosomes prohibiting stacking of U909 with the decoding nucleotide A919 (A1493 in *E. coli*). mtRF1a contacts the decoding nucleotides of the 12S rRNA via stacking interactions of V137 and R320 thereby stabilizing the switch loop in a similar conformation than in bacteria. The experimental EM density is shown as mesh. D) Superposition of mitochondrial ICT1 (ochre) and bacterial ArfB (violet, PDB = 6YSS (Chan et al., 2020)) upon alignment of the rRNA of the small subunit of the bacterial and mitochondrial ribosomes. The C-terminal α-helix is substantially longer in ICT1. Termination factors are shown for simplicity in isolation and tRNA^Met^ (green) and mRNA (red) of the ICT1 translation termination complex as well as adjacent ribosomal features (grey) are given for reference. DC = decoding center, PTC = peptidyl-transferase center E) Sequence conservation within the C-terminal domains of ICT1 and ArfB has been plotted as consensus sequence using weblogo (https://weblogo.berkeley.edu). Larger residues indicate a higher degree of conservation. Residues that make important interactions in the mitochondrial mRNA channel and are conserved in bacterial ArfB have been highlighted (asterisk). F – H) ICT1 binds the empty mRNA channel via its positively charged C-terminal extension. Interactions of the ICT1 C-terminal extension with the mRNA channel and the decoding nucleotides (green) or the tip of H69 (U909, purple) are shown with the corresponding experimental EM density as mesh. The 12S rRNA of the SSU is shown in yellow and residues conserved in bacterial ArfB are highlighted in light blue.

Binding of the stop codon by the codon recognition domain requires a rearrangement of the ribosomal decoding center, which occurs in a highly conserved fashion in bacteria and eukaryotes. There, the decoding nucleotide A1492 flips out of h44 while A1493 stacks with A1913 at the tip of helix 69 (H69) of the LSU 16S rRNA (*E. coli* numbering) (Fig. S1B). The tip of H69 is shortened in mitochondria lacking the otherwise conserved A1913 and contains U909 instead. Consequently, in the presence of the bulky codon recognition domain of mtRF1a, the tip of H69 is not able to bridge the distance to decoding nucleotide A919 (A1493 in *E. coli*). Instead, mitochondrial A919 engages in an alternative stacking interaction with Val137 of domain II of mtRF1a (Fig. 2C and S1B). In bacteria and eukaryotes, the second decoding nucleotide A918 (A1492 in *E. coli*) interacts with the switch loop of translation termination factors (Fig. S1B). Upon codon recognition, class I release factors such as mtRF1a that contain codon recognition and PTH domain typically undergo large-scale structural rearrangements to position the catalytic PTH domain into the PTC in order to induce peptidyl-tRNA hydrolysis.(Korostelev et al., 2008; Korostelev et al., 2010; Laurberg et al., 2008; Svidritskiy and Korostelev, 2018) The switch loop plays a critical role in coordinating codon recognition with structural rearrangement of the catalytic PTH domain. On the ribosome, the mtRF1a switch loop nestles into a pocket formed between uS12m, 12S rRNA and mtRF1a domain II (Fig. 2C). It adopts a conformation similar but not identical to bacterial RF1 with Arg320 stacking onto decoding nucleotide A918 (Fig. S1B).(Laurberg et al., 2008; Svidritskiy et al., 2016) Such staking interactions have been indicated to stabilize the repositioned PTH domain upon rearrangement of the decoding center in bacteria.(Korostelev et al., 2010; Svidritskiy and Korostelev, 2018; Svidritskiy et al., 2016) Intriguingly, removal of the stacking interaction as well as deletion of H69 have been shown to decrease the rate of translation termination more than 1000fold presumably due to diminished PTH accommodation.(Korostelev et al., 2010; Svidritskiy and Korostelev, 2018) Interestingly, a release factor mutant that shows less rigid accommodation of the PTH due to a truncation in the switch loop makes a more specific distinction between sense and stop codons albeit at a lower rate than wild type showing that the switch loop likely balances translation termination accuracy and efficiency in bacteria.(Svidritskiy and Korostelev, 2018) It will therefore be interesting to understand whether the rRNA variations in the mitochondrial decoding center lead to an altered discriminatory potential and different kinetics of mtRF1a binding to stop codons in comparison to bacteria.

### ICT1 recognizes a partially empty mRNA channel to rescue stalled ribosomes

Curiously, ICT1 is a mitoribosomal protein that is part of the large subunit close to the central protuberance (Fig. S2A and S2B).(Brown et al., 2014; Greber et al., 2014) However, it fulfills its function as translation termination factor as an extraribosomal copy.(Akabane et al., 2014; Greber et al., 2014) We solved the structure of ICT1 bound to the A site of a mitoribosome that contains a short hexanucleotide mRNA (5’CUGAUG3’) and mtRNA^Met^-fMet in the P site to a resolution of 3.4 Å (Fig. 1D). The short hexanucleotide leaves the mRNA channel partially empty mimicking a scenario, in which the ribosome is stalled on the 3’ end of an mRNA that lacks a stop codon. In this rescue complex, ICT1 places its conserved PTH domain with the catalytic GGQ motif close to the peptidyl transferase center in an orientation largely identical to mtRF1a. In contrast to mtRF1a, ICT1 lacks a codon recognition domain and has acquired a positively charged C-terminal extension instead. We find that this extension adopts a more than 40 Å long α-helical fold on the ribosome and inserts into the empty mRNA channel reaching from the decoding center towards the mRNA entry (Fig. S2B and S2C). Presence of an mRNA and the ICT1 C-terminal tail in the mRNA channel appear to be mutually exclusive, indicating that ICT1 specifically acts on mitoribosomes that contain truncated, aberrant mRNAs. This behavior is reminiscent of the bacterial rescue factor ArfB (YaeJ) that probes the occupancy of the mRNA channel in a similar way albeit its C-terminal α-helix is significantly shorter (Fig. 2D, Fig. S2C and S2D).(Carbone et al., 2020; Chan et al., 2020; Gagnon et al., 2012) ICT1 contains a number of amino acids in its C-terminal extension that are highly conserved with respect to bacterial ArfB (Fig. 2E). These residues have been shown to be functionally important since they are crucial for binding of ArfB to the bacterial ribosome and for PTH activity of ArfB as well as ICT1.(Kogure et al., 2014) We find in our model that the residues engage in extensive interactions with the SSU 12S rRNA deep inside the mRNA channel and at the decoding center as well as with uS12m - all ribosomal elements conserved from bacteria (Fig. 2F and 2G, Fig. S2E and S2F). Despite the similarity of the interactions with the bacterial system, we do also observe slight differences between binding modi of ICT1 and ArfB. For example, decoding nucleotide G256 stacks with Arg192 in ICT1 that is not conserved in bacterial ArfB to lift G256 into a tilted conformation (Fig. 2G and Fig.S2F). In addition, the extended alpha-helical element of ICT1 places His176 as a replacement for ArfB Pro110 for stacking interactions of the decoding nucleotide A919 and U909 at the tip of H69 and thereby compensates for evolutionary rRNA alterations in the mitoribosome (Fig. 2H and Fig. S2G). Taken together, our structural data explain how the C-terminal extension of ICT1 specifically associates with conserved rRNA elements of the vacant mRNA channel during ribosome rescue in a fashion reminiscent to bacterial ArfB. Moreover, our data show how mitochondria-specific alterations in ICT1 help to re-establish conserved ArfB interactions despite evolutionary changes in the mitoribosomal rRNA.

### Interaction of the catalytic PTH domain with the tRNA acceptor end

Upon recognition of the stop codon or the vacant mRNA channel, mtRF1a and ICT1 will trigger hydrolysis of the polypeptide chain from the tRNA acceptor (CCA) end. Hydrolysis occurs at the peptidyl transferase center, which guides the productive positioning of the catalytic GGQ motif of the peptidyl hydrolase domains of the termination factors. Consequently, the orientation of the PTH domains of mtRF1a and ICT1 towards the tRNA-acceptor (CCA) end is highly similar in both complexes (Fig. 3A). Because the PTH domain of ICT1 was substantially better resolved than the one from mtRF1a, we focus on its structural interpretation here. The catalytic ^88^GGQ^90^ motif is well visible and we find the side chain of the conserved glutamine Gln90 pointing away from the 3’ and 2’ OH group of the terminal adenosine A71 of tRNA^Met^ (Fig. 3B). This conformation is in line with a highly conserved mode of catalysis, in which the backbone NH group of Gln90 as well as the 3’ OH and 2’ OH group of A71 of peptidyl-tRNA are critical for cleavage of the ester bond between nascent chain and P site tRNA.(Rodnina, 2018) As our translation termination complexes contain wild-type termination factors, our EM density clearly corresponds to a posthydrolysis state, in which fMet has already been released from the 3’OH group of A71 of mtRNA^Met^.

**Fig 3.**
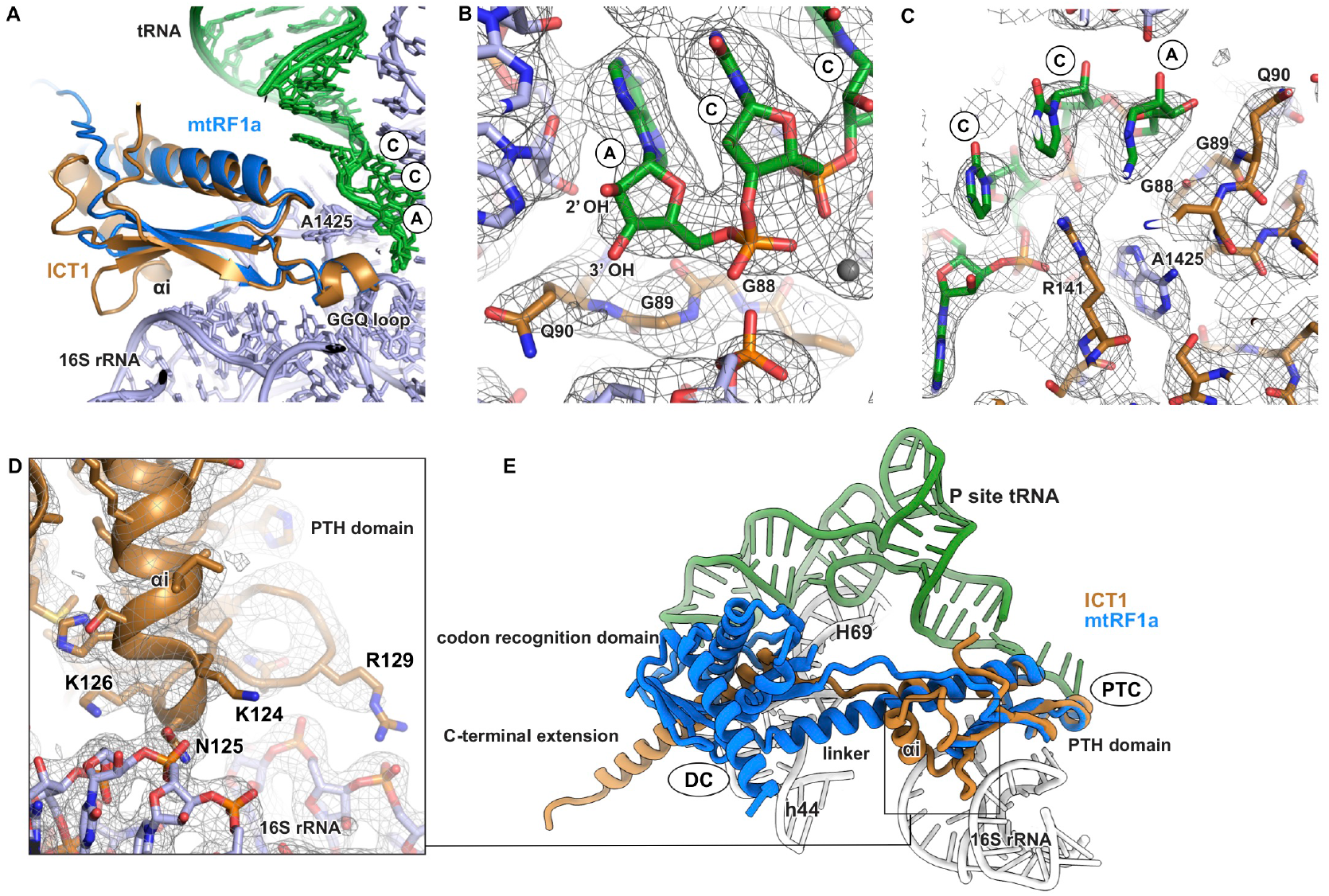
Positioning of the catalytic GGQ motif at the peptidyl-transferase center. A) The PTH domains of ICT1 (ochre) and mtRF1a (blue) are shown after superposition of the 16S rRNA of the mitoribosomal LSU. The highly conserved, catalytic GGQ motif is positioned at the peptidyl-transferase center (PTC) close to the tRNA acceptor (CCA) end (green). Binding of the ICT1 PTH domain is additionally stabilized by the 25 amino acid-long insertion αi. B) The catalytic GGQ motif of ICT1 and the tRNA CCA end are shown with the experimental EM density. The tRNA is deacylated as our complex contains wild type ICT1. The backbone NH group of Q90 and the 2’ OH and 3’ OH groups of the terminal ribose in the tRNA CCA end are crucial for hydrolysis of the ester bond between tRNA and nascent polypeptide chain. C) Positioning of the GGQ motif is aided by stacking of the highly conserved R141 onto rRNA nucleotide A1425. R141 in addition interacts with bases and phosphate backbone of the tRNA CCA end. D) Binding of the ICT1 PTH domain to the LSU is stabilized via a polypeptide insertion (αi) that is absent in mtRF1a. αi contacts the 16S rRNA of the LSU via multiple positively charged amino acid side chains (K124, K126, R129). The EM density is shown as mesh. E) Superposition of ICT1 (ochre) and mtRF1a (blue) after alignment of the 16S rRNA of the LSU. PTH domain and codon recognition domain of mtRF1a are connected via a rigid α-helix that may aid stable positioning of the PTH domain at the PTC. In ICT1, C-terminal extension and PTH domain are connected via a flexible linker and positioning of the PTH domain may require additional stabilization via αi. tRNA (green), h44 of the SSU, H69 of the LSU and the part of the 16S rRNA that interacts with αi of ICT1 are shown for reference (white). Important ribosomal sites are indicated. DC = decoding center, PTC = peptidyl-transferase center

Productive positioning of the catalytic GGQ motif depends on its interaction with the LSU 16S rRNA and the P site tRNA^Met^. A1425 (A2602 in *E. coli*) of the 16S rRNA serves as a critical anchoring point for correct GGQ loop placement into the PTC (Fig. 3C). A1425 is tightly embedded in the PTH domain by stacking interactions with a highly conserved arginine residue Arg141 in ICT1 that in addition contacts the phosphate backbone of the mtRNA^Met^ CCA end to fix the orientation of the GGQ motif close to the tRNA acceptor end (Fig. 3C). Notably, positioning of the PTH domain of ICT1 additionally depends on an α-helical, 25 amino acid insertion (termed αi) also found in bacterial ArfB but absent in mtRF1a (Fig. 3D and 3E).(Gagnon et al., 2012; Handa et al., 2010) Although αi appears not to play a role in ribosome binding, it was shown to be essential for catalytic activity of ICT1.(Akabane et al., 2014) In our complex, a patch of positively charged amino acid side chains (Lys124, Asn125, Lys126, Asn128, Arg129) in αi closely contact the phosphate backbone of 16S rRNA to stabilize association of the PTH domain with the LSU (Fig. 3D).(Akabane et al., 2014) Increased stability of the ICT1 PTH domain on the LSU may explain the higher quality of its EM density in comparison to our mtRF1a complex. A major difference between ICT1 and mtRF1a is that the orientation of the PTH domain of mtRF1a at the PTC is facilitated by a rigid α-helical connection between PTH and codon recognition domain. In contrast, the ICT1 C-terminal extension and the PTH domain are connected via a flexible linker, likely necessitating the additional interactions of αi with the LSU to establish a catalytically productive conformation of ICT1 (Fig. 3E).

### Architecture of the pre- and post-splitting mitoribosomal complexes

Once the nascent chain has been released from the ribosome, the ribosomal subunits are split in order to remove any remaining mRNA and tRNAs and to engage in the translation of the next mRNA. In order to recapitulate a mitochondrial recycling event, we isolated 55S mitoribosomes from porcine liver tissue. These mitoribosomes retain endogenous tRNA and mRNA molecules during purification. However, we realized that they usually carry tRNAs in their A and P sites making them a poor substrate for ribosome recycling, which requires a vacant A site. To establish a recycling-competent state of the mitoribosome, we therefore pre-treated our 55S mitoribosomes with the translation elongation factor mtEFG1 in the presence of GTP to promote translocation of the tRNA-mRNA module and clearance of the A site (Fig. 4A). After separation of 55S mitoribosomes from mtEFG1, we promoted ribosome recycling by the addition of mtRRF, mtEFG2 and mtIF3 in presence of the nonhydrolyzable nucleotide analogue GDPNP. In contrast to bacterial EFG, GTP binding to mtEFG2 but not hydrolysis is required for splitting of the mitoribosomal subunits.(Tsuboi et al., 2009) We included mtIF3 in the reaction as it has previously been suggested to prevent re-association of the mitoribosomal subunits and aid clearance of the tRNAs from the SSU.(Christian and Spremulli, 2009; Derbikova et al., 2018; Rudler et al., 2019)

**Fig 4.**
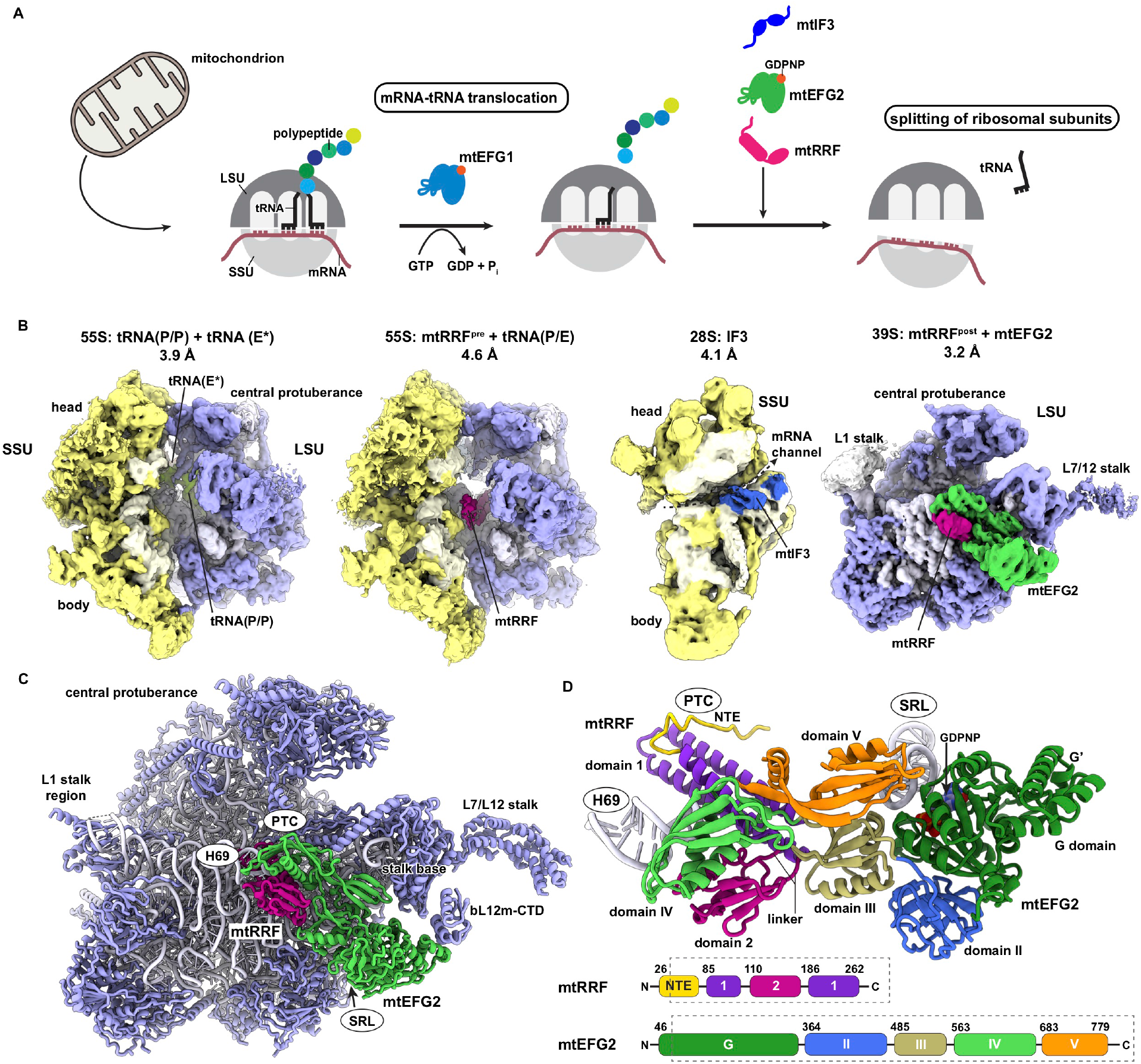
Architecture of the mitoribosomal recycling complex. A) Schematic illustration of the sample preparation strategy. Mitoribosomes were isolated from mitochondria and a recycling-competent state was derived by incubation with the translation elongation factor mtEFG1 in presence of GTP. Subsequently, mtRRF, mtEFG2 and mtIF3 were added to trigger dissociation of the ribosomal subunits. B) Ribosomal complexes contained in the recycling reaction are colour-coded according to SSU (yellow), and LSU (light blue). mtRRF (purple), mtIF3 (dark blue), and mtEFG2 (green) are shaded differently. A Gaussian filter (s.d. = 1.5) has been applied to the depicted EM densities. C) The structural model of the 39S post-splitting complex is shown with mtRRF in purple and mtEFG2 in green. mtEFG2 contacts the mitoribosome close to the L7/12 stalk base and interacts extensively with mtRRF next to 16S rRNA helix 69 (H69). mtEFG2 is in addition contacted by the bL12m-CTD of the L7/12 stalk. D) The complex of mtEFG2 and mtRRF complex is shown in isolation and adjacent ribosomal elements are labeled (white). Protein domains are colour-coded and indicated in the schematics below. Numbers indicate amino acid residues at the domain boundaries. PTC = peptidyl-transferase center, SRL= sarcin ricin loop

Cryo-EM analysis of the mitoribosome recycling reaction unraveled four distinct complexes that we could classify as pre- and post-splitting states (Fig. 4B). In the pre-splitting state, we can identify two distinct assemblies: the 55S mitoribosome containing mtRRF and a tRNA bound in the hybrid state between ribosomal P and E site (termed 55S: mtRRF + tRNA(P/E)), and the 55S monosome with an E site tRNA bound in a non-canonical position (termed 55S: tRNA(P/P) + tRNA(E*)). In the post-splitting state, we find mtIF3 weakly associated to the 28S SSU (termed 28S: IF3), and the 39S LSU in complex with mtRRF and mtEFG2 (termed 39S: mtRRF + mtEFG2). As mtIF3 bound to the 28S has been visualized previously (Khawaja et al., 2020; Koripella et al., 2019a), we did not interpret it here and concentrated on obtaining structural models for both pre-splitting complexes and the 39S:mtRRF + mtEFG2 post-splitting complex. In 39S: mtRRF + mtEFG2, both recycling factors extensively interact with one another and with the LSU subunit interface (Fig. 3C). Here, mtRRF blocks the peptidyl-transferase center (PTC) and nestles around H69 of the LSU 16S rRNA. mtEFG2 exhibits an overall domain architecture and position on the LSU similar to the translation elongation factor mtEFG1 (Fig. 4C and 4D, Fig. S3C and 3D).(Koripella et al., 2020b; Kummer and Ban, 2020) This includes contacts of the mtEFG2 G domain with the sarcin ricin loop (SRL), and of domain V with the ribosomal stalk base. Moreover, we find the C-terminal domain of bL12m (bL12m-CTD) engaged with the GTPase (G) domain of mtEFG2. bL12m-CTD plays an important role in recruitment of translational GTPases to the ribosome and has previously been shown to also contact the G domains of mitochondrial translation factors mtIF2 and mtEFG1.(Koripella et al., 2020b; Kummer and Ban, 2020; Kummer et al., 2018)

Despite lacking any translation factors, one of the pre-splitting complexes provides intriguing insights into a novel and likely mitochondria-specific mode of tRNA binding. In 55S: tRNA(P/P) + tRNA(E*), we identify two tRNA molecules, one of which adopts a canonical P/P position, whereas the other is bound to the ribosomal E site in a highly unusual conformation, which we term tRNA(E*) here. It shows a twisted and almost perpendicular orientation with respect to the canonical E/E state (Fig. S3A and S3B). The CCA end of tRNA(E*) maintains its interaction with the canonical binding pocket on the LSU, whereas its anticodon stem loop (ASL) has escaped from the canonical E site position on the SSU and rather engages with a mitoribosome-specific rRNA insertion (A739-C747) between helices 41 and 42 of 12S rRNA (Fig. S3A). This tRNA conformation is not only distinct from the canonical A, P, and E positions but also from the more recently described Z-site tRNA on the cytosolic ribosome.(Brown et al., 2018) Although the mammalian mitoribosome has generally undergone a reduction of its rRNA core, the A739-C747 RNA segment represents a rare case of rRNA expansion. The mitoribosomal tRNA binding sites as well as the L1 stalk, which plays a critical role in ejection of deacylated tRNAs from the mitoribosome, have been extensively remodeled with respect to the bacterial ancestor.(Amunts et al., 2015; Greber et al., 2015) We propose that the identified short rRNA expansion segment may be one of the new features mediating binding and movement of tRNAs on the mitoribosome and that remodeling of tRNA-binding sites may be accompanied by alternative modes of tRNA binding and ejection at the mitoribosomal E site.

### Concerted action of mtRRF and mtEFG2 splits the mitoribosomal subunits

As our EM dataset contains pre- and post-splitting recycling complexes, it allows us to understand the conformational rearrangements that trigger mitoribosomal subunit dissociation. In the pre-splitting 55S: mtRRF + tRNA(P/E), we visualize how mtRRF interacts with the mitoribosome prior to mtEFG2 binding (Fig. 5A). The overall structure of mtRRF is similar to its bacterial counterpart with the factor composed of two domains (Fig 4D).(Agrawal et al., 2004; Borovinskaya et al., 2007; Gao et al., 2005; Koripella et al., 2019b; Selmer et al., 1999; Weixlbaumer et al., 2007; Wilson et al., 2005) Domain 1 is a triple helix bundle that intimately engages with the phosphate backbone of the LSU 16S rRNA via numerous positively charged amino acid side chains to stabilize binding of the factor to the ribosome (Fig. 4D and Fig. S3E). A linker region connects domain 1 with domain 2 that is flexibly disposed with weak contacts to uS12m on the SSU (Fig. 5A). Location of mtRRF domain 1 across A and P sites of the LSU blocks the PTC for engagement with tRNA (Fig. S3F). Instead, the bound tRNA adopts a hybrid position (tRNA(P/E)), in which the ASL is still bound to the P site on the SSU but the acceptor arm has already moved towards the E site (Fig. 5A). Only a tRNA that is deacylated can adopt such a conformation, indicating that - in analogy to the bacterial system - mitoribosomal recycling depends on prior hydrolysis of the peptidyl bond to ensure that recycling can only occur on mitoribosomes that finished translation.(Agrawal et al., 2004; Dunkle et al., 2011; Fu et al., 2016; Zhou et al., 2020) Mammalian mtRRF carries a mitochondria-specific N-terminal extension, which we are able to partially resolve here. This extension forms a hydrophobic cluster, in which the ribosomal protein uL16m inserts a polypeptide loop via an apical methionine residue Met136 (Fig. 5B). Flipping of uL16m has not been shown in earlier reports on mtRRF engaged with the mitoribosome possibly due to lower local resolution in these studies.(Aibara et al., 2020; Koripella et al., 2019b) The hydrophobic pocket in the mtRRF NTE and the hydrophobic residue at the tip of the uS16m are conserved in mammalian mitochondria indicating that flipping of uL16m may occur more generally and may provide an additional measure to stabilize mtRRF binding on the LSU and to trigger repositioning of deacylated P-site tRNA into a hybrid P/E position (Fig. 5C). Besides this well-resolved hydrophobic pocket, the remaining part of the NTE remains invisible probably due to structural flexibility. Recently, it has been proposed that the N-terminal extension contains a small subdomain deposited between SSU 12S rRNA h44, LSU 16S rRNA H69 and mtRRF domain 1 to stabilize a rotated conformation of the mitoribosome that promotes mtEFG2 binding to the recycling complex.(Koripella et al., 2020a) Interestingly, this subdomain appears to encompass residues 1-11 of the NTE, which are consistently attributed by prediction algorithms to be part of the cleavable mitochondrial import sequence (MTS) of mtRRF. In contrast to this study, we do not use full-length mtRRF here but a construct lacking the predicted MTS of 26 amino acids at the N-terminus. Although the MTS has initially been speculated to be retained during import of mtRRF into mitochondria, it was shown not to be required for ribosome binding.(Rorbach et al., 2008) In accordance, we observe efficient ribosome recycling in the absence of the predicted MTS as the majority of our initial 55S particles has been split into 39S and 28S subpopulations in the EM dataset (Fig. S9). This indicates that the N-terminal 26 amino acids of mtRRF are at least *in vitro* not critical for recycling of the mitoribosome.

**Fig 5.**
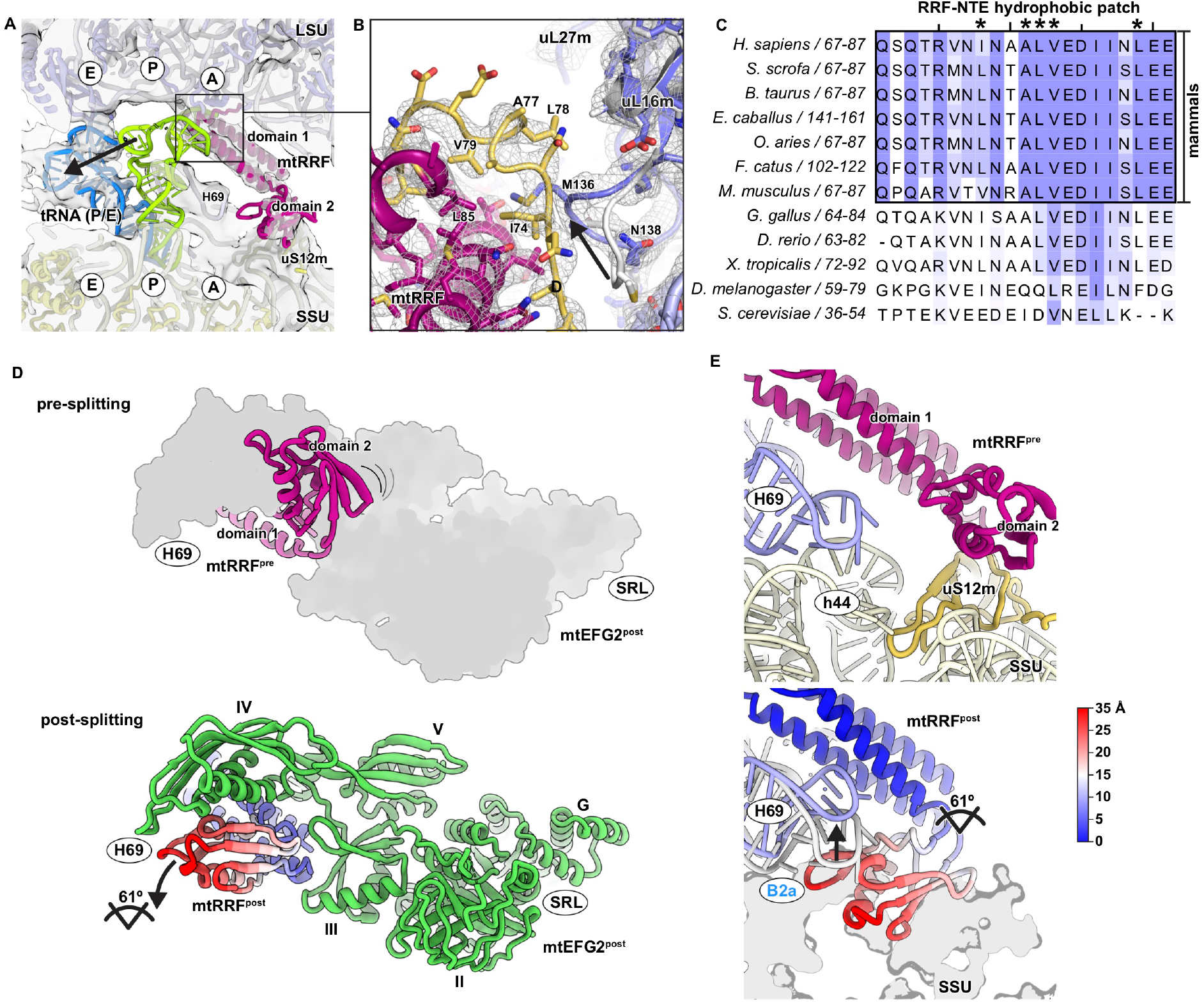
Comparison of pre- and post-splitting complexes shows mechanism of subunit dissociation. A) The subunit interface of the pre-splitting mitoribosome with mtRRF (purple) and tRNA in the hybrid (P/E) position (blue) is depicted with the corresponding Gaussian-filtered EM density (s.d. = 1.5). A tRNA in P/P position (green) has been fitted for reference and ribosomal tRNA binding sites on the LSU (light blue) and SSU (yellow) are indicated by the letters A, P, and E. B) mtRRF carries a mitochondria-specific N-terminal extension (yellow) that forms a hydrophobic pocket, into which a loop of uL16m of the LSU inserts an apical hydrophobic side chain. The experimental EM density is shown as mesh. C) The hydrophobic pocket in the mtRRF N-terminal extension (NTE) is conserved among mammals. NTE sequences have been aligned using Clustal Omega and coloured according to conservation in Jalview (conservation threshold = 30 overall, and 50 for the mammalian NTE) (Waterhouse et al., 2009). Residues that form the hydrophobic pocket are labeled with asterisk. D) Domain 2 of mtRRF undergoes a large-scale rotational motion upon mtEFG2 binding to split the ribosomal subunits. Pre-(top) and post-splitting (bottom) conformation of mtRRF (mtRRF^pre^ in purple) are shown in context of mtEFG2. mtRRF would clash with mtEFG2 (shown as grey outline) in its conformation adopted in the pre-splitting complex. However, mtRRF domain 2 is bend into a conformation compatible with mtEFG2 (green) association in the post-splitting complex. mtRRF^post^ has been coloured according to the displacement of the atoms with respect to mtRRF^pre^ showing that domain 1 retains its position on the LSU whereas domain 2 undergoes a large rotational movement of about 61 degree. The corresponding colour key for mtRRF^post^ is identical to the one shown in Fig. 5E. Rotation axis and angle have been calculated in PyMOL using the draw_rotation_axis.py script (P.G. Calvo). Adjacent ribosomal sites are indicated (SRL = sarcin ricin loop). E) Motion of mtRRF domain 2 contributes to dissolution to the B2a intersubunit bridge. In the pre-state (top), domain 2 is loosely associated to uS12m and 16S rRNA helix 69 (H69) of the LSU and 12S rRNA helix 44 (h44) of the SSU interact to form the intersubunit bridge B2a. In the post-splitting complex (bottom), H69 motion disrupts B2a and mtRRF domain 2 clashes with the SSU 12S rRNA (h44) leading to dissociation of the ribosomal subunits. The SSU is shown as a grey outline and the positions of H69 in the pre-splitting (white) and in the post-splitting complex (blue) is depicted. mtRRF^post^ is colour-coded according to displacement of the atoms with respect to mtRRF^pre^. The corresponding colour key is given.

Comparison of 55S: mtRRF + tRNA(P/E) with the post-splitting complex 39S: mtRRF + mtEFG2 provides important indications how concerted action of mtRRF and mtEFG2 may break the 55S mitoribosome apart. Binding of mtEFG2 to the ribosome causes a large-scale rotational motion of mtRRF domain 2 of about 61 degree around its linker region while domain 1 stably retains its position on the 16S rRNA (Fig. 5D and 5E). An analogous conformational rearrangement of the recycling factor occurs in bacteria upon binding EFG.(Fu et al., 2016; Gao et al., 2007; Gao et al., 2005; Zhou et al., 2020) mtRRF domain 2 movement is critical to avoid clashes with mtEFG2 domain IV, which appears to directly push mtRRF into its alternative conformation (Fig. 5D). The positioning of mtRRF domain 2 and mtEFG4 domain IV in the post-splitting complex is sterically incompatible with the presence of the 12S SSU rRNA h44 when the two subunits are associated in the 55S mitoribosome. Moreover, H69 of 16S LSU rRNA becomes tightly embedded in mtRRF domain 1 and 2, which lifts it away from the interaction site with h44 (Fig. 5E and Fig. S4A). Concerted action of mtRRF and mtEFG2 thereby disrupts the critical mitoribosomal intersubunit bridge B2a, eventually resulting in dissociation of the ribosomal subunits (Fig. 5E and Fig. S4B).(Zhang et al., 2015) In fact, H69 is not only shifted by interaction with mtRRF but in addition experiences a more general reorganization by alterations in its base stacking (Fig. S4C). Moreover, U912 of H69 flips outwards to insert in between U958 and U960 of the underlying H71 thereby possibly stabilizing the compressed H69 conformation during recycling.

### mtEFG2 is specialized for ribosome recycling via a lock-and-key mechanism

Mitochondria contain two paralogues of the canonical translation elongation factor found in many bacteria with distinct functions in polypeptide synthesis. While mtEFG1 catalyzes tRNA-mRNA movement during translation elongation, mtEFG2 is dedicated to ribosome recycling.(Tsuboi et al., 2009) Interestingly, also a number of bacteria harbor two versions of the elongation factor indicating that the mitochondrial principle of task division might be more broadly used. Our data now reveal how this functional specialization is established. As the primary sequences of mtEFG1 and mtEFG2 are only 33% identical, we started with a more general analysis of the surface properties of both proteins. Plotting of the electrostatic potential on the molecular surface shows that especially in regions where mtEFG2 contacts mtRRF during ribosome recycling, mtEFG1 and mtEFG2 are vastly different (Fig. 6A and 6B). This is especially prominent considering the contacts of mtEFG2 with the linker region of mtRRF that is critical for rotational motion of mtRRF domain 2 and consequently subunit splitting (Fig. 6B and 6C). In case of mtEFG2 and mtRRF, interacting surfaces electrostatically match while the contact site in mtEFG1 contains a number of positively charged amino acid side chains that would be repulsive for mtRRF binding (Fig. 6C, Fig. S4D and S4E). The second prominent discrepancy between mtRRF and mtEFG1 occurs due to a mitochondria-specific C-terminal extension of mtEFG1, which is absent in mtEFG2. Part of this extension forms a rigid α-helix that would sterically clash with mtRRF domain 1 and thus precludes mtEFG1 from contributing to ribosome recycling (Fig. 6D and 6E). In summary, mtEFG2 is electrostatically and sterically optimized for interaction with mtRRF while mtEFG1 has acquired a range of features preventing its involvement in ribosome recycling.

**Fig 6.**
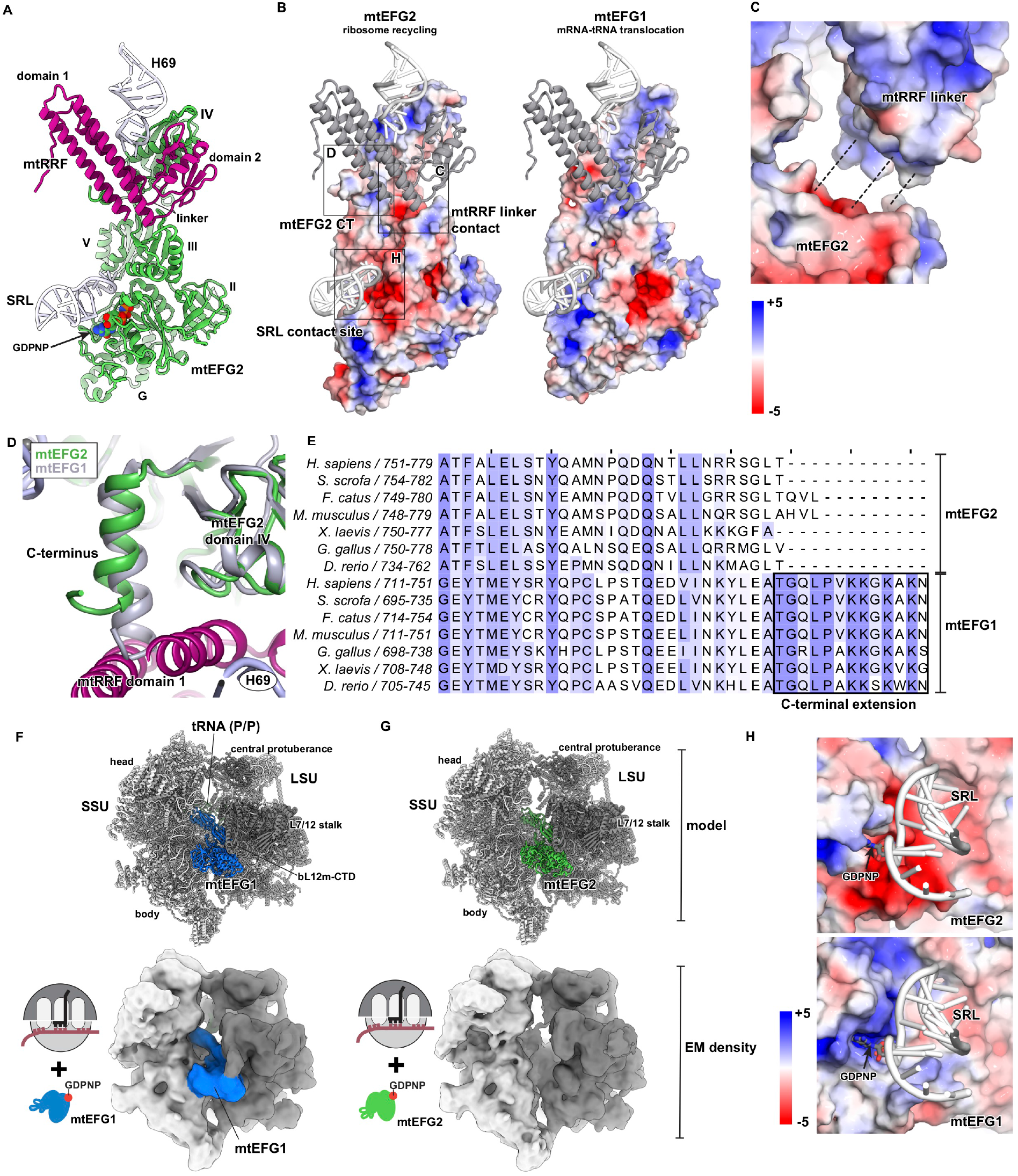
mtEFG2 is specialized for ribosome recycling. A) Overview of the interaction of mtEFG2 (green) and mtRRF (purple) with adjacent ribosomal rRNA elements shown for reference (white). SRL = sarcin ricin loop B) mtEFG2 electrostatically matches mtRRF whereas mtEFG1 does not. The electrostatic potential of mtEFG1 and mtEFG2 has been calculated in PyMOL using the APBS tool (Baker et al., 2001) and plotted on the surface of both proteins. The scale bar shows the corresponding colour code between ± 5 kT/e. mtRRF is shown in grey. Positioning of mtRRF onto mtEFG1 was done via alignment with mtEFG2. Boxes highlight regions of the mtEFG2 surface that show prominent differences with respect to mtEFG1. Letters inside the boxes refer to the corresponding panel of Fig.6, in which the boxed area is shown as close-up. CT = C-terminus C) A closer view on the interaction area between the mtRRF linker region and the corresponding surface on mtEFG2 indicates favourable electrostatic interactions. The surfaces of both proteins are displayed adjacent but not in direct contact to allow inspection of the interaction area. Attracting interactions between the two surfaces are indicated with dashed lines. D) mtEFG1 carries a C-terminal extension that is sterically incompatible with binding to mtRRF (purple). mtEFG1 (grey) has been superpositioned onto mtEFG2 (green). Each domain of mtEFG1 has been aligned separately to account for slightly distinct conformations of mtEFG1 and mtEFG2 on the ribosome. The mtEFG1 C-terminal extension has been fitted together with domain IV with which it makes prominent contacts. The C-terminal extension clearly clashes with domain 1 of mtRRF in this conformation. E) Clustal Omega sequence alignment for the C-terminal regions of mtEFG2 and mtEFG1 have been coloured according to conservation in Jalview. The overall conservation threshold was set to 30 and for the boxed area containing the mtEFG1-specific C-terminal extension to 50. Species and amino acid borders of the aligned sequences are given on the left. F – G) 55S mitoribosomes have been prepared as indicated in Fig. 4A and incubated with mtEFG1 (F) or mtEFG2 (G), respectively, in presence of the non-hydrolyzable nucleotide analogue GDPNP. While mtEFG1 is able to bind the mitoribosome under these conditions, mtEFG2 is not. On top, structural models are depicted to indicate the location of mtEFG1 and mtEFG2 on the ribosome. Ribosomal subunits are shown in grey, mtEFG1 in blue, tRNA in dark green and mtEFG2 in light green. In case of mtEFG1, PDB 6YDP was used for display (Kummer and Ban, 2020). For mtEFG2, our 39S: mtRRF + mtEFG2 model was superpositioned onto the 55S mitoribosome of PDB 6YDP via the 16S rRNA to show the predicted location of mtEFG2 on the 55S mitoribosome. At the bottom, the experimentally derived EM densities for both reactions are coloured according to the structural models on top. The EM density in (F) accounts for mtEFG1 but in case of mtEFG2 the corresponding density is missing (G). H) Electrostatic surface potential calculated in PyMOL using the APBS tool and plotted on the molecular surface of mtEFG2 and mtEFG1, respectively, at a range between ± 5 kT/e. The images are close-ups of the binding region of both proteins to the SRL. In mtEFG2, this region shows a strong negative surface potential indicating repulsive interactions with the phosphate backbone of the sarcin ricin loop (SRL).

### mtEFG2 is a poor ribosome binder and requires mtRRF for selective targeting to the mitoribosome

Our data also reveal that specialization of mitochondrial elongation factors may not only arise due to optimized interaction surfaces but may already be established at the step of translation factor recruitment to the ribosome. During our reconstitution approaches, we consistently observed that mtEFG2 is unable to bind the 55S mitoribosome in the absence of mtRRF, even when the non-hydrolyzable nucleotide analogues GDPNP or GTPgS are added (Fig. 6G and Fig. S11). However, when mtRRF is added to the reaction, mtEFG2 efficiently engages the ribosomal complex to trigger recycling. Our data therefore suggest that mtEFG2 alone may be a poor ribosome binder and appears to depend on mtRRF for recruitment to the mitoribosome. This is in stark contrast to mtEFG1, which can engage the mitoribosome efficiently on its own in the presence of GDPNP (Fig. 6F and Fig. S10).(Koripella et al., 2020b; Kummer and Ban, 2020)

Several features of mtEFG2 may explain its limited ability to bind the ribosome independently. One is a strikingly negative surface potential of the region of its G domain that interacts with the phosphate backbone of the SRL. A negative electrostatic surface would predict a repulsive instead of an attracting interaction between the mitoribosomal rRNA and mtEFG2 (Fig. 6H). In addition, we find that the density for bL12m-CTD, which facilitates binding of translational GTPases to the GTPase-associated center (GAC) of the ribosome, is rather weak in our 39S: mtRRF + mtEFG2 complex suggesting that bL12m-CTD is less stably associated to mtEFG2 than to mtEFG1 (Fig. S5A and S5B). In addition, bL12m-CTD adopts an unusual, rotated conformation on the G’ domain of mtEFG2 probably caused by a mtEF2-specific polypeptide insertion in G’ (Fig. S5D and S5E). We have shown earlier that closure of the GAC onto mtEFG1 may stabilize it in a translocation-competent conformation on the ribosome.(Kummer and Ban, 2020) Although the GAC moves upon mtEFG2 binding to the mitoribosome, this motion follows a distinct trajectory probably due to a slightly distinct position of mtEFG2 domain V on the LSU in the recycling complex (Fig. S5F and S5G). In this conformation, mtEFG2 appears to engage in less stabilizing contacts with the GAC than mtEFG1 (Fig. S5G). Overall, several indications suggest that mtEFG2 properties may limit its interaction with the mitoribosome and that it may require mtRRF for stable ribosome engagement. Functionally, these observations have important consequences. The dependence on mtRRF may prevent competitive binding of mtEFG2 to the elongating mitoribosome, where it could hamper mtEFG1 action and slow down translation elongation. Moreover, it ensures that mtRRF binds the ribosome prior to mtEFG2 and thereby establishes an order of events during ribosome recycling. First the peptidyl-tRNA bond needs to be cleaved to allow tRNA relocation and mtRRF association, and only subsequently mtEFG2 binding triggers splitting of the ribosomal subunits.

## DISCUSSION

Mitochondria were predicted to contain four distinct translation termination factors. Here, we used an extensive EM screening approach to clarify the contribution of these factors to translation termination in mammalian mitochondria. In this screen, we find mtRF1a in association with the canonical stop codons UAA and UAG on the mitoribosome in accordance with previous biochemical data.(Nozaki et al., 2008; Soleimanpour-Lichaei et al., 2007) We show that mtRF1a decodes canonical stop codons using structural features that are surprisingly conserved considering the striking evolutionary divergence of the mitochondrial translation system (Fig. 7). Additionally, we show that the ICT1 acts as a ribosome rescue factor that specifically associates with ribosomes with a partially empty mRNA channel in line with an earlier biochemical study (Fig. 7).(Feaga et al., 2016) Such a scenario typically arises when mitoribosomes stall on aberrant, truncated messages and the lack of a canonical stop codon prevents mtRF1a action. We did not find release factor candidates mtRF1 and C12ORF65 bound to the mitochondrial ribosome in any of the conditions tested. This may either indicate that they rather fulfill functions outside of the ribosome or that they may require additional interaction partners for ribosome binding that have not yet been identified. In fact, C12ORF65 has only very recently been shown to cooperate with the RNA-binding protein MTRES1 (C6ORF203) in an alternative mitoribosome rescue pathway that acts on the large subunit only.(Desai et al., 2020) In combination with our data, this shows that despite similar domain architecture and related electrostatic properties, ICT1 and C12ORF65 follow distinct modes of action to recover stalled mitoribosomes for translation and it may also indicate that their interaction with the mitoribosome could be controlled by different cues. While ICT1 appears to detect truncated mRNAs on the mitoribosome, the initial trigger that commits stalled mitoribosomes for rescue by C12ORF65 binding has not been established. Interestingly, ICT1 has more recently emerged as an important oncogene associated with unfavourable prognosis and was shown to be upregulated in a variety of cancer types including breast cancer, prostate cancer, lung cancer, and leukemia.(Chang et al., 2017; Huang et al., 2014; Lao et al., 2016; Peng et al., 2018; Tao et al., 2017; Wang et al., 2017; Xie et al., 2015) Our structural study may therefore be helpful for the development of novel therapeutic approaches to counteract malignancy and promote cancer regression. Intriguingly, none of the four termination factors directly recognized the alternative stop codons in our screen. This may, again, be caused by the absence of critical interaction partners in our *in vitro* setup or may indicate that a currently still unidentified factor or mechanism may promote translation termination on alternative human stop codons AGA and AGG. Previously, bioinformatic and biochemical studies have implied ICT1 or mtRF1 in recognition of alternative stop codons. We however did not find compelling evidence for these suggestions in our setup. (Akabane et al., 2014; Lind et al., 2013; Richter et al., 2010; Young et al., 2010) Alternatively, a -1 frameshifting has been proposed to enable termination at these sites by placing a canonical stop codon in the ribosomal A site.(Temperley et al., 2010) As there is currently no *in vitro* translation system available for mitochondria, we have not been able to conclusively test this hypothesis here. A third putative mechanism to terminate translation at alternative stop codons could involve the recently identified ribosome rescue pathway by C12ORF65 and MTRES1. However, here the factors that could recognize the terminating ribosome and split the ribosomal subunits prior to action of the two rescue factors still have to be identified.

**Fig 7.**
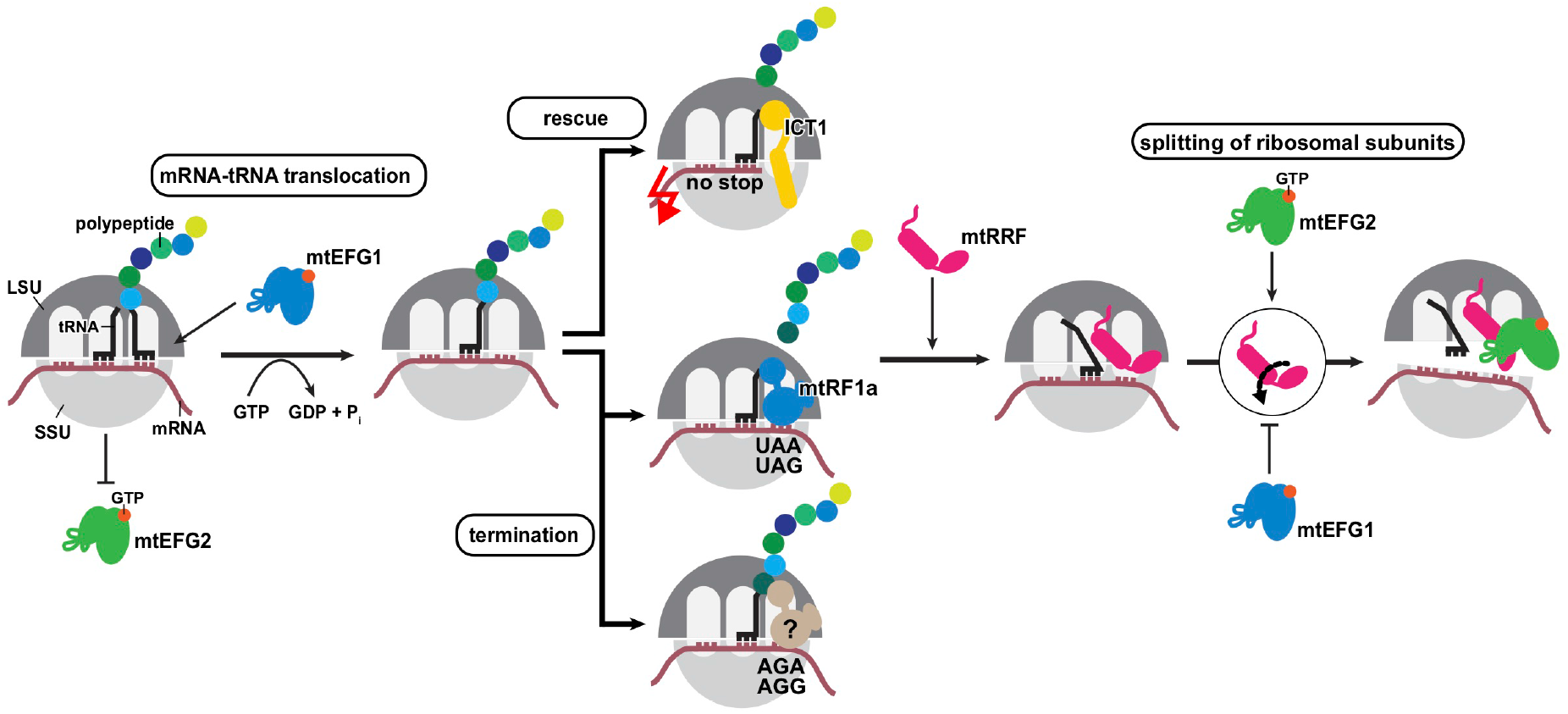
Model of mitochondrial translation termination and ribosome recycling. Mitochondria contain two elongation factors termed mtEFG1 and mtEFG2. mtEFG1 is specialized for the process of translation elongation where it catalyzes the movement of the tRNA and mRNA through the ribosome, whereas mtEFG2 is specialized for ribosome recycling. Polypeptide synthesis is terminated when the ribosome encounters the end of the open reading frame and a stop codon is placed in the ribosomal A site, which can be one of the canonical stop codons (UAA, UAG) or a non-canonical, mitochondria-specific stop codon (AGA, AGG). Canonical stop codons are recognized by the translation termination factor mtRF1a via highly conserved recognition motifs. Non-canonical stop codons appear not to be recognized – at least directly – by any of the predicted, mitochondrial translation termination factors. When a stop codon is recognized, the termination factor catalyzes the release of the nascent chain from the peptidyl-tRNA. The nascent chain gets also released, when the mitoribosome stalls on messages that are aberrant and do not contain a stop codon. In this case, the mitochondrial rescue factor ICT1 binds to the partially empty mRNA channel on the SSU and triggers peptidyl hydrolysis. Once the nascent chain has been released, ribosome recycling occurs in an ordered series of events. First, mtRRF binds to the vacant A site of the ribosome and induces the P site tRNA to adopt a hybrid P/E conformation. mtRRF acts as an affinity anchor for subsequent recruitment of mtEFG2, whose binding leads to a large-scale rotational motion of mtRRF domain 2. In a concerted action, both factors cause dissociation of the mitoribosomal subunits by dissolution of the intersubunit bridge B2a and steric interference with the SSU. Specialization of both elongation factors is caused by distinct properties of both proteins. mtEFG1 is electrostatically and sterically imcompatible with mtRRF thereby preventing its participation in ribosome recycling. mtEFG2 shows surface properties that make it a poor ribosome binder on its own necessitating the presence of mtRRF for ribosome binding. Consequently, mtEFG2 is unable to interfere with mtEFG1 during translation elongation ensuring efficient polypeptide synthesis.

In addition to our insights on nascent chain release by mtRF1a at canonical stop codons and ribosome rescue by ICT1, we visualize how the concerted action of mtRRF and mtEFG2 recycles the mitoribosome after polypeptide hydrolysis for the next round of translation. We show that large-scale rotational motions of mtRRF induced by binding of mtEFG2 cause subunit dissociation by dissolution of the mitoribosomal intersubunit bridge B2a and by steric interference of both recycling factors with the 12S rRNA of the SSU (Fig. 7). Our data clarify that mitochondrial mtEFG2 has specialized for its interaction with mtRRF electrostatically and sterically whereas the second elongation factor mtEFG1 shows properties that prevent its participation in ribosome recycling (Fig. 7). We thereby establish the molecular basis how mtEFG1 and mtEFG2 have specialized to fulfill distinct tasks during mitochondrial protein synthesis. Additionally, we propose a novel mechanism, in which mtEFG2 is a poor ribosome binder and its recruitment depends on prior association of mtRRF to the mitoribosome (Fig. 7). Such a mode of action has two consequences: it prevents interference of mtEFG2 with mtEFG1 action during translation elongation, and it establishes an order of events that leads to productive ribosome recycling.

## Acknowledgements

We thank A. Filipovska and M. Leibundgut for critical reading of the manuscript. We are grateful to D. Boehringer and S. Mattei for support during EM data acquisition as well as the ScopeM staff for technical support. E.K. was supported by an EMBO long-term fellowship (1196-2014). K.N.S. was supported by a Ph.D. fellowship by Böhringer Ingelheim Fonds. This work was supported by the Swiss National Science Foundation grant (310030B_163478), via the National Centre of Excellence in RNA and Disease and project funding 138262 as well as an ETH Research Grant ETH-23 18-2 to N.B.

## Author contribution

E.K., K.N.S. and N.B. conceptualized the project. E.K., K.N.S. and T.S. prepared the samples. E.K. and K.N.S. collected the EM data for the translation termination factor screening at the F20 and analyzed them. E.K., K.N.S. and A.S. collected the high-resolution EM data for the translation termination and ribosome recycling complexes at the Krios and E.K. and K.N.S. analyzed them. E.K. built and validated the structural models with technical support from A.S. E.K., K.N.S. and N.B. interpreted the structural data. E.K. wrote the manuscript and prepared the figures. All authors contributed to the editing of the manuscript into its final form.

## Data availability

Structural models and cryo-EM maps have been deposited in the Protein Data Bank (PDB) and Electron Microscopy Data Bank (EMDB), respectively under following accession codes. PDB 7NQH and EMD-12527 for 55S: mtRNA^Met^ + mtRF1a, PDB 7NQL and EMD-12529 for 55S: mtRNA^Met^ + ICT1, PDB 7NSH and EMD-12567 for 39S: mtRRF + mtEFG2, PDB 7NSI and EMD-12568 for 55S: mtRRF + tRNA(P/E), and PDB 7NSJ and EMD-12569 for 55S: tRNA(P/P) + tRNA(E*).

## Methods

### Plasmids

Open reading frames (ORFs) for MT-CO3(AGA) and MT-CO3(UAA) were generated from the previously used MT-CO3 construct by site-directed mutagenesis introducing a stop codon directly following the AUG start codon (Kummer et al., 2018). Human mtRF1a, mtRF1, ICT1, and C12ORF65 were ordered from GenScript, and the open reading frames for mtEFG1, mtEFG2 and mtRRF were ordered from Thermo Scientific. All ORFs were purchased codon-optimized and including a N-terminal His_6_-tag, a TEV cleavage site and a GGSGSG linker. The ORFs were subcloned into pET24a for recombinant protein production in *E. coli*.

### Preparation of human mitochondrial translation factors

Mitochondrial translation initiation factors mtIF2 and mtIF3 were purified as described earlier (Kummer et al., 2018).

The constructs for all mitochondrial translation termination factors (mtRF1, mtRF1a, ICT1, and C12ORF65) were expressed from pET24a and contained an N-terminal His_6_-tag followed by a TEV cleavage site and a GGSGSG linker. Expression constructs were designed such that they did not contain the predicted N-terminal mitochondrial localization sequences (MitoFates and MitoProtII were used for MTS prediction). Specifically, mtRF1a lacked 32 amino acids, C12orf65 lacked 25 amino acids, and ICT1 lacked 29 amino acids at their N-termini, respectively. mtRF1 was initially expressed as construct lacking 57 N-terminal amino acids. However this construct has heavily aggregation-prone and we therefore worked with a slightly shorter, but soluble version of mtRF1 that did not contain the first 74 amino acids. His_6_-mtRF1a was expressed in *E. coli* BL21 SI pRARE at 18 °C overnight and purified using a HisTrap FF 5-ml column (GE Healthcare) in standard buffers (50 mM HEPES-KOH pH 7.6, 100 mM KCl, 5 mM MgCl_2_, 40 or 500 mM imidazole, 10% (w/v) glycerol, 0.5 mM TCEP). His_6_-tagged TEV protease was added to the protein and the sample was incubated at 4 °C overnight. Another HisTrap FF 5-ml column purification followed to remove the TEV-protease, uncleaved protein and the His_6_-tag. Cleaved protein was applied to a desalting column 26/10 (GE Healthcare) and thereby buffer exchanged to the storage buffer (40 mM HEPES-KOH pH 7.6, 200 mM KCl, 40 mM MgCl_2_, 10% (w/v) glycerol, 0.5 mM TCEP).

His_6_-mtRF1 was expressed in BL21 (DE3) that contained in addition the chaperone-encoding plasmid pG-KJE8 (Takara Bio Inc.). The cells were cultured in the presence of 5 ng/μl tetracycline and 0.5 mg/ml L-arabinose to induce chaperone expression. Protein production was carried out overnight at 18 °C. mtRF1 was purified using a 5 ml HisTrap HP column coupled to a 5 ml Heparin column in standard buffers (50 mM HEPES-KOH pH 7.0, 100 or 800 mM KCl, 20 mM MgCl_2_, 40 or 500 mM imidazole, 10% (w/v) glycerol, 1 mM DTT) and the His_6_-tag was removed as described before. The cleaved protein was further purified on a Superdex200 16/600 and at the same time buffer exchanged into storage buffer (20 mM HEPES-KOH pH 7.0, 150 mM KCl, 20 mM MgCl_2_, 40 or 500 mM imidazole, 10% (w/v) glycerol, 1 mM DTT). His_6_-ICT1 and His_6_-C12ORF65 were expressed in *E. coli* BL21 (DE3) pLysS at 30 °C for 3 h and purified as described for mtRF1a using a HisTrap FF 5-ml column. In case of C12ORF65, the HisTrap FF column was additionally coupled to a HiTrap Heparin HP 5-ml column (GE Healthcare). Both proteins were subjected to TEV protease cleavage at 4 °C overnight and another HisTrap FF 5-ml column purification step. In case of ICT1, the protein was subjected to size-exclusion chromatography on a HighLoad 16/60 Superdex75 (GE Healthcare).

All proteins were buffer exchanged, when required, and concentrated in the storage buffer (40 mM HEPES-KOH pH 7.6, 200 mM KCl, 40 mM MgCl_2_, 10% (w/v) glycerol, 1 mM TCEP) using an Amicon Ultra-15 centrifugal filter (30-kDa molecular weight cut-off), aliquoted, flash-frozen and stored at -80 °C until further use.

mtEFG1, mtEFG2, and mtRRF were expressed in *E. coli* BL21 (DE3) at 18°C overnight. mtEFG1 and mtEFG2 were isolated via affinity purification on HisTrap FF 5 ml columns (GE Healthcare) and standard buffers as described above. The His_6_-tag was cleaved by incubation with TEV-His_6_ protease overnight at 4°C. Uncleaved protein, the His_6_-tag, and TEV-His_6_ protease were removed by reverse Ni^2+^-based affinity chromatography using a HisTrap FF 5 ml column. Finally, mtEFG1 and mtEFG2 were buffer exchanged into storage buffer (40 mM HEPES-KOH pH 7.6, 200 mM KCl, 40 mM MgCl_2_, 1 mM DTT, 10% (w/v) glycerol) by size exclusion chromatography using a Superdex200 16/600 column (GE Healthcare). Aliquots were flash frozen in liquid nitrogen and stored at −80°C until further use. mtRRF was purified using HisTrap FF 5ml and HiTrap Heparin HP 5ml columns (GE Healthcare) in tandem with standard buffers as described for mtRF1. The His_6_-tag was removed by TEV-cleavage overnight at 4°C and the cleaved protein was separated from the His_6_ tag and His_6_-TEV protease via reverse His-affinity purification on a HisTrap FF 5 ml. mtRRF was exchanged into storage buffer (50 mM HEPES-KOH pH 7.6, 150 mM KCl, 20 mM MgCl_2_, 10 % (w/v) glycerol, 1 mM DTT) and concentrated using an Amicon Ultra 15 (molecular weight cutoff at 10 kDa). Aliquots were flash frozen in liquid nitrogen and stored at -80 °C until further use.

### Preparation of human mitochondrial mRNAs

MT-CO3(AGA), MT-CO3(UAA) and MT-CO3(UAG) were designed as hammerhead - MT-CO3 fusion constructs in a pUC19 vector under control of a T7 promoter. The DNA templates were digested with StyI and subsequently purified by phenol-chloroform extraction and ethanol precipitation. CO3 mRNAs of approximately 200 nucleotide in length were generated from these linearized, purified MT-CO3 genes by *in vitro* run-off transcription. Transcription reactions (40 mM Tris-HCl pH 7.8, 30 mM MgCl_2_, 0.01% (v/v) Triton X-100, 5 mM DTT, 1 mM spermidine, 10 mM NTPs) were performed at 37 °C overnight and transcripts were afterwards incubated for 5 min at 95 °C, then for 5 min on ice, again 5 min at 95 °C and 15 min on ice to maximize hammerhead cleavage. Preparative urea PAGE (5% polyacrylamide, 1x TBE, 7 M urea) was performed to purify the cleaved mRNA transcripts from the hammerhead ribozyme. The respective mRNA bands were excised and the mRNAs were extracted by shaking of the gel pieces in water at 4 °C overnight. Finally, the mRNAs were buffer exchanged in an Amicon Ultra-15 centrifugal filter into water to remove any residual urea (10-kDa molecular weight cut-off), aliquoted and stored at - 20 °C until further use. The short RNA oligonucleotide with the sequence CUGAUG employed for forming the no-stop translation termination complex has been ordered from Dharmacon.

### Preparation of mitochondrial fMet-tRNA^Met^

Formylated and aminoacylated fMet-tRNA^Met^ was produced following published protocols (Kummer et al., 2018). In brief, mitochondrial tRNA^Met^ was generated via run-off T7 transcription and hammerhead ribozyme cleavage. The RNA fragment was purified via agarose gel electrophoresis, folded in the presence of 10 mM MgCl_2_, aminoacylated and formylated *in vitro*, and purified via phenol-chloroform extraction. Aliquots were flash frozen and stored at −80°C.

### Preparation of mitoribosomal subunits and the pre-splitting 55S mitoribosome

Mitochondria were isolated from porcine liver tissue as described earlier (Greber et al., 2014) and stored at -80°C until further use. Mitoribosomal subunits for reconstitution of translation termination complexes were purified from mitochondria following protocols established earlier (Kummer et al., 2018). 55S mitoribosomes for ribosome recycling reactions were isolated similarly to the mitoribosomal subunits. In brief, mitochondria were lysed in 1x MB (20 mM HEPES-KOH pH 7.6, 100 mM KCl, 40 mM MgCl_2_, and 1 mM DTT) by thawing and dounce homogenizing in presence of RNase inhibitor (RiboLock, Thermo Scientific). Subsequently, mitoribosomes were solubilized from the mitochondrial membranes by addition of 1.6 % (v/v) Triton X-100 for 30 min at 4°C. After two short clearance spins (Type 45 Ti rotor from Beckman Coulter, 15 min at 20000 rpm), lysates were spun for 24 h at 4°C at 50000 rpm in a Type 70 Ti rotor (Beckman Coulter) through a 40 % (w/v) sucrose cushion in 1x MB. Mitoribosomal pellets were resuspended by gently shaking in 1x MB for 60 min on ice. 55S mitoribosomes then underwent a clearance spin at 14000 rpm and 4°C (tabletop centrifuge 5427 R, Eppendorf). We realized that our mitoribosomes contained endogeneous tRNAs in the A and P sites of the ribosome. As ribosome recycling requires a vacant A site, we incubated the 55S mitoribosomes with 10 μM mtEFG1 and 5 mM GTP to induce translocation of the A site tRNA. mtEFG1 was subsequently removed from the mitoribosomes via a 10 – 40 % (w/v) sucrose gradient in 1x MB (17 h, 22500 rpm, SW 32 Ti rotor (Beckman Coulter), 4 °C). The gradients were fractionated and mitoribosome-containing fraction were pooled. The sucrose-containing buffer was exchanged against 1x MB using an Amicon Ultra 0.5 (molecular weight cutoff at 100 kDa). Concentrated and buffer-exchanged 55S mitoribosomes were then used directly for preparation of the recycling complex.

### Preparation of translation termination and ribosome recycling complexes

Translation termination complexes were generated starting with the assembly of the translation initiation complex from mitoribosomal subunits. This was done to ensure efficient and correct positioning of mRNA and mtRNA^Met^-fMet in the P site of the ribosome. The initiation complex was assembled as described earlier (Kummer et al., 2018), except for using GTP instead of GTPγS to allow mtIF2 to dissociate from the ribosome leaving a vacant A-site. To achieve the presence of canonical (UAA, UAG) and non-canonical (AGA) stop codons in the ribosomal A site, mitochondrial initiation complexes were formed in the presence of MT-CO3 mRNA containing the respective stop codon right after the AUG start codon. In case of the no-stop scenario, in which the ribosome stalls on the 3’ end of the mRNA, a short RNA oligo (CUGAUG, Dharmacon) was used as mRNA instead. The putative release factors mtRF1a, mtRF1, ICT1 or C12ORF65 (final concentration 1 μM) were added after incubating the initiation complex for 5 min at 37 °C. After another 3 min at 37 °C the termination complexes were placed on ice for 15 min and subsequently applied to Quantifoil R2/2 holey carbon grids (Quantifoil Micro Tools) either coated with a thin continuous carbon film or with graphene oxide flakes (Sigma-Aldrich). The grids were vitrified by plunge-freezing in pure ethane on a Vitrobot (FEI).

Ribosome recycling complexes were assembled from 55S mitoribosomes pre-treated with mtEFG1 that contained endogeneous mRNAs and tRNAs. We realized that our 55S mitoribosomes did not have to be treated with puromycin for efficient ribosome recycling indicating that the nascent chain has been released from the peptidyl-tRNA already during preparation of the mitoribosomes. Puromycin was therefore omitted from the reaction. 60 nM of 55S mitoribosomes were incubated with 1 μM mtRRF, 1 μM mtIF3 and 2 mM GDPNP for 3 min at 37 °C and 1 μM of mtEFG2 was added afterwards. The recycling reaction was left for 3 more min at 37 °C and then placed on ice before applying the sample to Quantifoil R2/2 holey carbon grids covered with a thin continuous carbon film. The sample was finally plunge-frozen on a Vitrobot IV (FEI) in a liquid ethane/propane mixture.

### Cryo-EM based release and recycling factor screening and image processing

Screening of translation termination complexes and of ribosome recycling complexes containing mtEFG1 and mtEFG2 in presence of GDPNP was carried out at a FEI Tecnai F20 equipped with a Falcon II direct electron detector using EPU. Images were collected at 82400x magnification and a total dose of 40 e^-^/Å^2^ either as integrated images or movies depending on the required exposure time. Movies were aligned, summed and dose-weighted in MOTIONCOR2 (Grant and Grigorieff, 2015; Zheng et al., 2017). For translation termination complexes, CTF estimation was done in GCTF (Zhang, 2016) and particle picking was carried out using the reference-free method based on the Laplacian-of-Gaussian (LoG) filter in RELION3 (Scheres, 2012; Zivanov et al., 2018).

In case of translation termination complexes, RELION3 was also used for the following cryo-EM reconstruction, in which reassembled 55S mitoribosomes were enriched. Local classification of the ribosomal A site was applied in a final step to discriminate empty ribosomes from the putative population of release factor bound ribosomes. In case of mitoribosomes incubated with mtEFG1 and mtEFG2 in presence of GDPNP, EM data were analyzed using cryoSPARC (Punjani et al., 2017) including patch CTF estimation, particle picking by the blob picker (minimum particle diameter 200 Å, maximum particle diameter 350 Å) and 2D classification. After 2D classification for 30 online EM iterations using 100 classes, particles from classes containing clear density for the 55S or ribosomal subunits were selected and underwent further classification by 3D heterogeneous refinement. As initial reference volume we used EMD-2914 (Greber et al., 2015) with the density for tRNAs subtracted. Classes with intact 55S mitoribosomes were pooled and underwent another round of 3D homogeneous refinement in cryoSPARC before display.

### Krios data collection of translation termination and ribosome recycling complexes

Images were acquired in movie mode on a FEI Titan Krios cryo-electron microscope using a Falcon III direct electron detector (FEI) at an accelerator voltage of 300 kV and a total electron dose of 40 e^-^/Å^2^. Images were collected at 100,719x magnification (pixel size of 1.39 Å/pixel) and a defocus range from -1.2 to 2.4 μm in case of translation termination factors mtRF1a and ICT1. Images of the ribosome recycling sample were collected at a smaller pixel size (1.087 Å/pixel) at 129,000x magnification. Movie frames were aligned, summed and dose-weighted in MOTIONCOR2 using 5×5 patches.(Grant and Grigorieff, 2015; Zheng et al., 2017) CTF estimation and particle selection were done using GCTF and BATCHBOXER (Ludtke et al., 1999; Zhang, 2016). After manual inspection of the micrographs, particles were picked using projections of the 55S mitoribosome as a reference (excluding tRNAs in A and P site) (PDB 5AJ4 (Greber et al., 2015)).

In case of translation termination complexes, data processing was performed in RELION3. 4x binned particle images were subjected to initial reference-free 2D classification and 3D classifications using the 55S mitoribosome as initial reference volume. Classes containing properly reassembled 55S mitoribosomes were joined and further refined and classified in 3D in RELION3 as indicated in the classification schemes (Fig. S7 and Fig. S8) to enrich the population of release factor-bound mitochondrial ribosomes. After re-extraction of unbinned particle images and final gold standard 3D refinement and postprocessing in RELION3, maps were used for model building.

In case of the ribosome-recycling sample, data were analyzed by combining elements from cryoSPARC with elements from RELION. First, particles were picked using Batchboxer with the 55S mitoribosome as picking reference. Of note, particles picked using the 55S mitoribosome or the 39S LSU did not appear to deviate and we thus decided to use the 55S as a picking reference. 4x binned particle images were extracted in Relion3 and imported into cryoSPARC for 2D classification using 40 online EM iterations and 200 classes. Subsequently, 3D heterogeneous refinement was performed against 6 initial reference volumes, two of which represented the 55S mitoribosome, two the 39S LSU and two the 28S SSU. The resulting 3D reconstructions contained one class of 28S SSU with mtIF3 bound, which was not further analyzed. Another class contained intact 55S particles that were re-extracted as unbinned images in RELION3 and imported into cryoSPARC. Here, the particle images were further classified by 3D heterogeneous refinement separating them into mitoribosomes containing mtRRF on the one hand and mitoribosomes containing tRNA (P/P) on the other hand. Both particle subsets were transferred to Relion3 for further 3D classifications without angular searches, involving local classification for tRNA(E*) in case of the 55S mitoribsome with tRNA(P/P) or classification of the entire particle in case of 55S mitoribosomes containing mtRRF to remove remaining poor particle images. Cleaned and classified particle populations were subjected to final 3D gold-standard refinement and postprocessed in RELION3. Resolutions of the 3D reconstructions were first estimated in Relion3 using FSC weighting. For the final EM maps used for model building, FSC weighting was however skipped and the maps were ad hoc filtered to the previously estimated resolutions to improve the overall map quality and the cross-correlation during subsequent real-space refinement in PHENIX. The third class extracted from the initial 3D heterogeneous refinement in cryoSPARC contained the 39S LSU in complex with mtRRF and mtEFG2. These particles were re-extracted as unbinned images in RELION3 and imported into cryoSPARC, where they were further classified by heterogeneous refinement to remove 39S particles from the population that did not contain the recycling factors. As the 3D heterogeneous refinement did not result in substantial separation of distinct 39S populations, we shifted the particle images selected from the best class of the heterogeneous refinement back to RELION3. After aligning the images in a gold-standard 3D refinement, 3D classification without angular searches was done for the entire particle. This led to separation of poor 39S subunits and 39S subunits that did not contain the recycling factors from a 39S class containing mtRRF and mtEFG2. Further local 3D classification using a mask around both recycling factors did not improve particle sorting or the density of the reconstruction and was thus abandoned. The particles underwent final gold-standard 3D refinement and postprocessing including FSC weighting in RELION3 and the resulting map was used for structure building and refinement.

### Structure building and refinement

Structure building was started by fitting the large ribosomal subunit, the head and body of the small ribosomal subunit, and the mtRNA^Met^ as well as homology models for the respective translations factors separately into the EM maps in Chimera. The structures of the mitochondrial translation initiation complex and the elongation POST complex served as starting points for the mitoribosome and the mtRNA^Met^ (PDBs 6GAW (Kummer et al., 2018) and 6YDP (Kummer and Ban, 2020)). Homology models of translation factors were generated in Phyre2 using PDB 4V63 as template for mtRF1a (Laurberg et al., 2008), 6YDP for mtEFG2 (Kummer and Ban, 2020), and 1EK8 (Kim et al., 2000)for mtRRF. In case of ICT1, its mitoribosomal copy from PDB 6GAW was taken as starting model and residues of human ICT1 that are divergent from pig were mutated to the human sequence in COOT (Emsley et al., 2010). Manual model building in COOT and O (Jones, 2004) was done to revise mitoribosomal features and associated factors.

In case of mtRF1a and ICT1, manual model building was followed by an initial round of reciprocal space refinement in PHENIX.REFINE to locate EM density that has is not accounted for by the structural models (Adams et al., 2010). Difference density maps were used for further manual model revision in COOT and O followed by a final real-space refinement in PHENIX using default restraints (Ramachandran plot, C-beta deviations, rotamer, secondary structure) and global minimization and B-factor refinement for 3 to 5 iterations with a weight between experimental data and restraints of 1.0 (Afonine et al., 2018).

For the recycling complexes, models were also manually adjusted in COOT after initial docking of rigid bodies into the EM maps in Chimera. The higher resolved 39S: mtRRF + mtEFG2 complex was used for manual revision of the recycling factors. In case of both 55S reconstructions, we relied mostly on docking of rigid bodies with mild manual adjustments, as the resolution did not permit near-atomic model building. Adjustments included linker regions of the rRNA between SSU head and body to reconnect both parts of the SSU. For 55S: mtRRF + tRNA(P/E), mtRRF domain 1 and domain 2 have been fitted separately as rigid bodies and the connecting linker was adjusted manually. Although we observe density for tRNAs in the A, P and E tRNA binding sites of the mitoribosome in this complex, the A site density was rather weak and we thus excluded it from interpretation during model building. For tRNA(P/E), the anticodon stem loop and the acceptor arm of the tRNA^Met^ were fitted as separate rigid bodies and manually reconnected to yield the hybrid tRNA conformation. The tRNAs in 55S: tRNA(P/P) and tRNA(E*) were initially fitted using the complete tRNA^Met^ (P/P) from PDB 6YDP (Kummer and Ban, 2020) as tRNA(P/P) or the acceptor arm and anticodon stem loop as separate entities that were reconnected to build tRNA(E*). For the mRNA in 55S: tRNA(P/P) + tRNA(E*), PDB 7A5I (Desai et al., 2020) was used as template. The sequences of tRNA^Met^ and the corresponding mRNAs were retained during real-space refinement and afterwards mutated to poly-uridine stretches to indicate that these RNAs resemble a mixture of endogenous ribonucleotides. Model building was followed by real-space refinement in PHENIX using defaults restraints (Ramachandran, C-beta deviations, rotamer, secondary structure) and global minimization as well as B factor refinement for 5 iterations with a weight of experimental data and restraints set to 1.0. PHENIX model validation statistics are shown in Table S1. Moreover, FSCs of the half sets of the experimental data as well as of the full map with the structural model, local resolution estimations, and angular particle distribution are included in Fig. S6.

## Figure generation

Molecular graphics were generated using the PyMOL (Schroedinger), UCSF Chimera or ChimeraX packages (Goddard et al., 2018; Pettersen et al., 2004).

