## Supplementary Material for "Release, rescue and recycling: termination of translation in mammalian mitochondria"

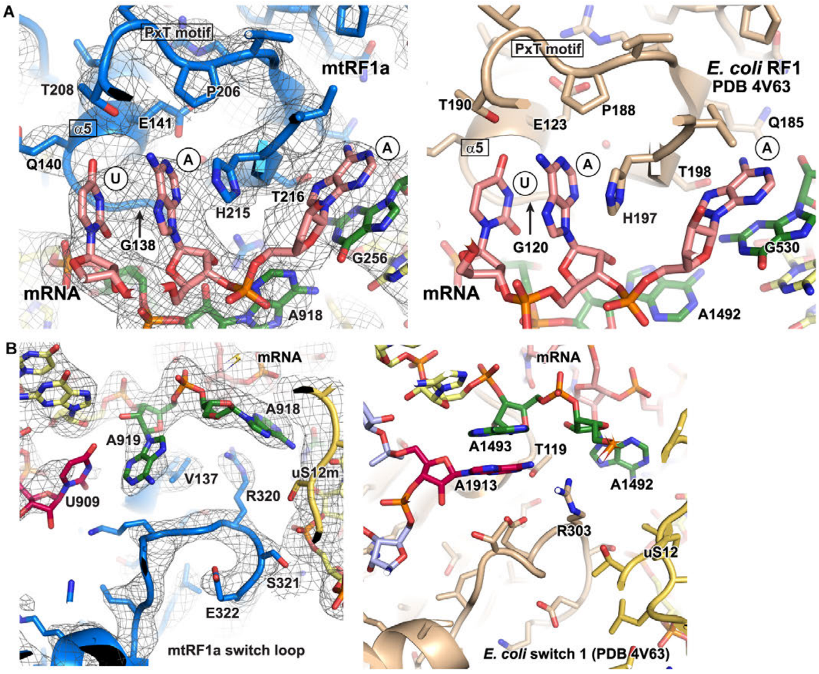


**Figure S1** Comparison of mitochondrial and bacterial translation termination factors, Related to Figure 2

A) Codon recognition motifs of mtRF1a (blue) and the bacterial RF1 (beige, PDB = 4V63 (Laurberg et al., 2008)) are shown visualizing their conservation. The UAA stop codon of the mRNA (salmon) and decoding nucleotides (green) of the ribosomal RNA of the SSU are highlighted. Experimental EM density for mtRF1a is shown as mesh.

B) Interaction of the switch loop in mtRF1a and bacterial RF1 (beige, PDB = 4V63 (Laurberg et al., 2008)) with the decoding center and ribosomal protein S12 (yellow) is displayed. Decoding bases are highlighted in green and the apical residue of H69 in purple. In contrast to bacteria, the tip of mitoribosomal H69 is too short to allow stacking of U909 onto decoding nucleotide A919. Experimental EM density for mtRF1a is shown as mesh.

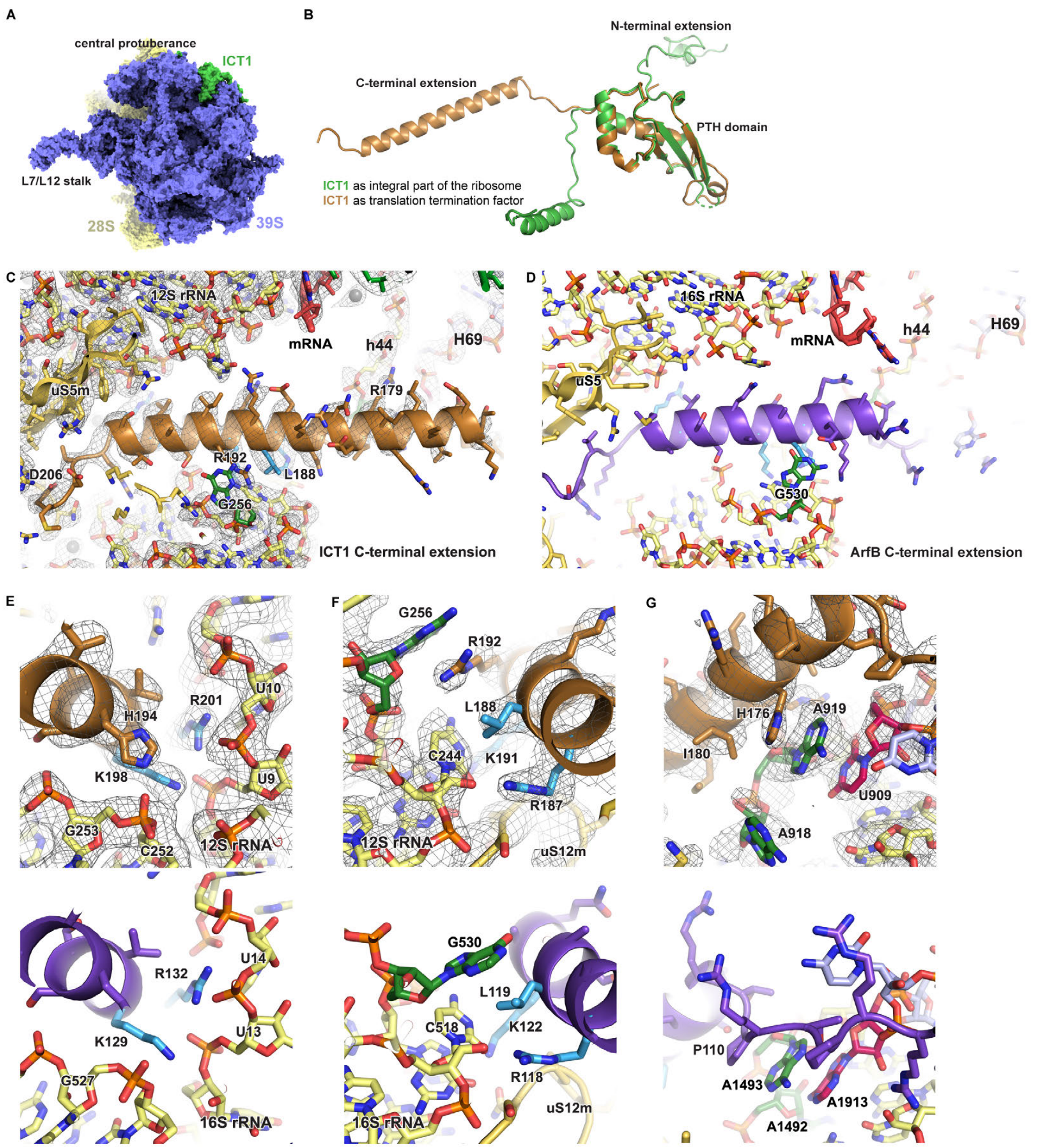


**Figure S2** Comparison of ICT1 to the bacterial homolog ArfB, Related to Figure 2

A) The structural model of the mitochondrial ribosome is shown in a side view as surface representation with the SSU colored yellow and the LSU blue. ICT1 in green is a mitoribosomal protein of the LSU located close to the central protuberance.

B) Superposition of ICT1 as a ribosomal protein (green) and in its extra-ribosomal conformation as translation termination factor (ochre). The structure of the PTH domain is retained while the C-terminal extension adopts distinct conformations.

C – D) The C-terminal extension of ICT1 and bacterial ArfB (violet, PDB = 6YSS (Chan et al., 2020)) are shown in the context of the ribosome. The extension is substantially longer in ICT1 but some of the residues that interact with the ribosomal RNA in the mRNA channel of the SSU are highly conserved (light blue). Decoding nucleotides are colored green and the experimental EM density for ICT1 is shown as mesh.

E – G) ICT1 binds the empty mRNA channel via its positively charged C-terminal extension. On top, interactions of the ICT1 C-terminal extension with the mRNA channel and the decoding nucleotides (green) or the tip of H69 (U909, purple) are shown with the corresponding experimental EM density as mesh. The 12S rRNA (mitochondria) is shown in yellow and residues conserved in bacterial ArfB are highlighted in light blue. At the bottom, identical views of the bacterial ribosome in complex with ArfB are shown for reference (PDB = 6YSS (Chan et al., 2020)).

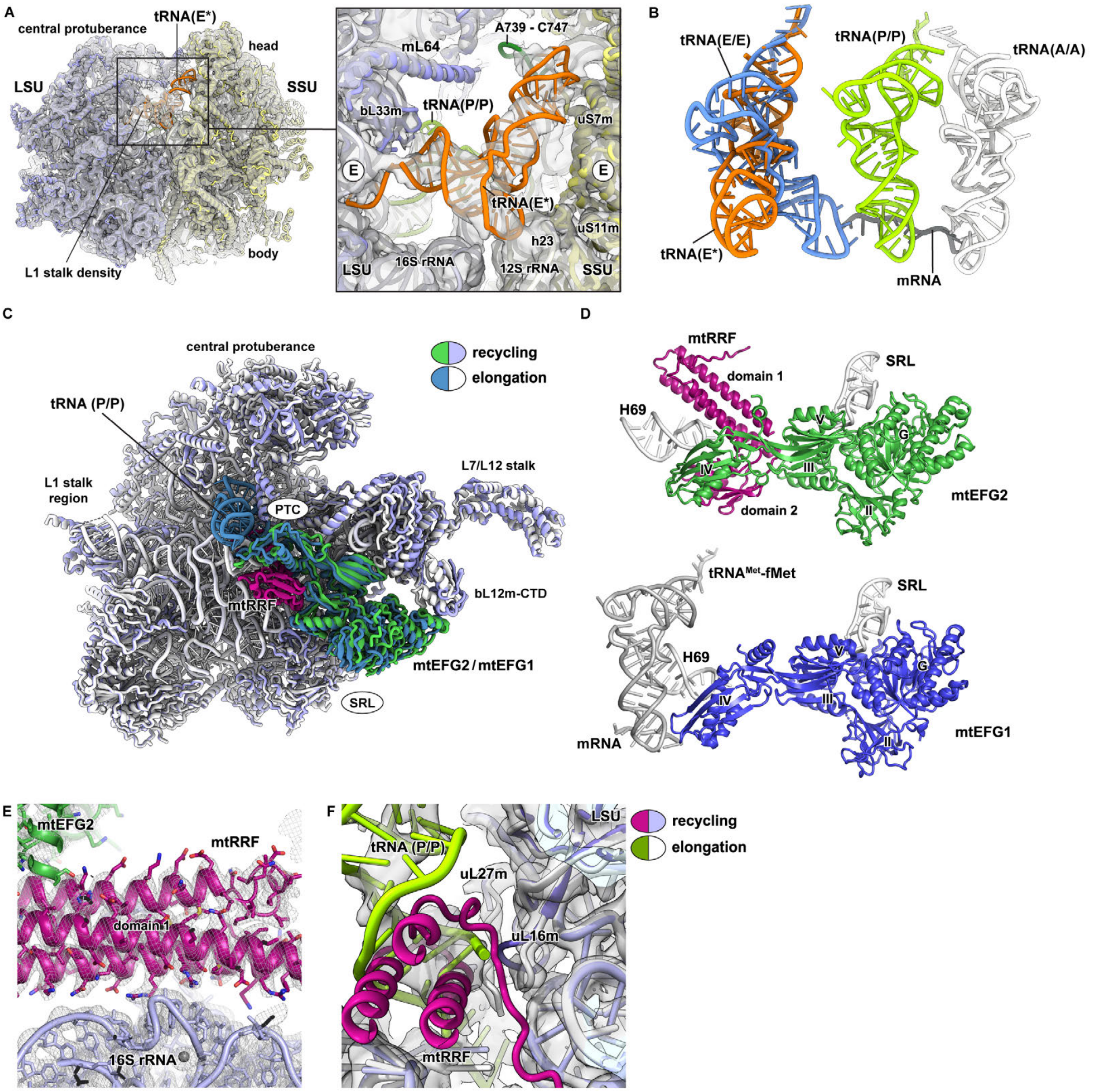


**Figure S3** Architecture of ribosome recycling complexes, Related to Figure 4 and Figure 5

A) The 55S ribosome in complex with a tRNA in the P/P position and an unconventionally bound E site tRNA, which we term tRNA(E*) here. The EM density has been Gaussian-filtered (s.d. = 1.5) and the underlying structural model is colored according to LSU (blue), SSU (yellow), tRNA(P/P) (green), and tRNA(E*) (orange). Details of the tRNA(E*) interaction with the ribosome are shown on the right. While the CCA end retains its position in the canonical E site on the LSU, the anticodon stem loop slips out of its binding site on the SSU and interacts with a mitochondria-specific rRNA insertion (A739-C747) that is highlighted in dark green. The tRNA elbow contacts h23 of the 12S SSU rRNA leading to an overall conformation of tRNA(E*) on the ribosome that is almost perpendicular to a classical E site tRNA.

B) tRNA(E*) is shown in relation to the classical positions of tRNAs in the A (white), P (green), and E (blue) sites of the ribosome. tRNA(P/P) and tRNA(E*) models are derived from this study. tRNA(A/A) and tRNA(E/E) are shown after superposition of the 12S rRNA of the SSU of our structural model with PDB 7A5I (Desai et al., 2020).

C – D) Overview of a superposition of the translation elongation complex (LSU in white, mtEFG1 and P site tRNA in blue) with the ribosome recycling complex (LSU in light blue, mtRRF in purple, mtEFG2 in green). mtEFG1 and mtEFG2 are shown in the context of the LSU and in isolation. Both elongation factors adopt similar conformations and positions on the ribosome. The bL12m C-terminal domain (bL12m-CTD) interacts with the GTPase (G) domains of both factors to aid their recruitment to the ribosome. SRL = sarcin ricin loop, PTC = peptidyl transferase center

E) Binding of mtRRF domain 1 (purple) to the 16S rRNA (light blue) of the LSU is achieved via interaction of numerous positively charged amino acid side chains with the phosphate backbone of the ribosomal RNA.

F) Superposition of the elongation complex (LSU in white and P site tRNA in green, PDB = 6YDP (Kummer and Ban, 2020)) with the ribosome recycling complex (LSU in light blue and mtRRF in purple) indicates that mtRRF binding to the ribosome is incompatible with the P/P position of a peptidyl-tRNA at the ribosomal peptidyl transferase center. The corresponding experimental EM density for PDB 6YDP is shown as semi-transparent surface (EMD-10778).

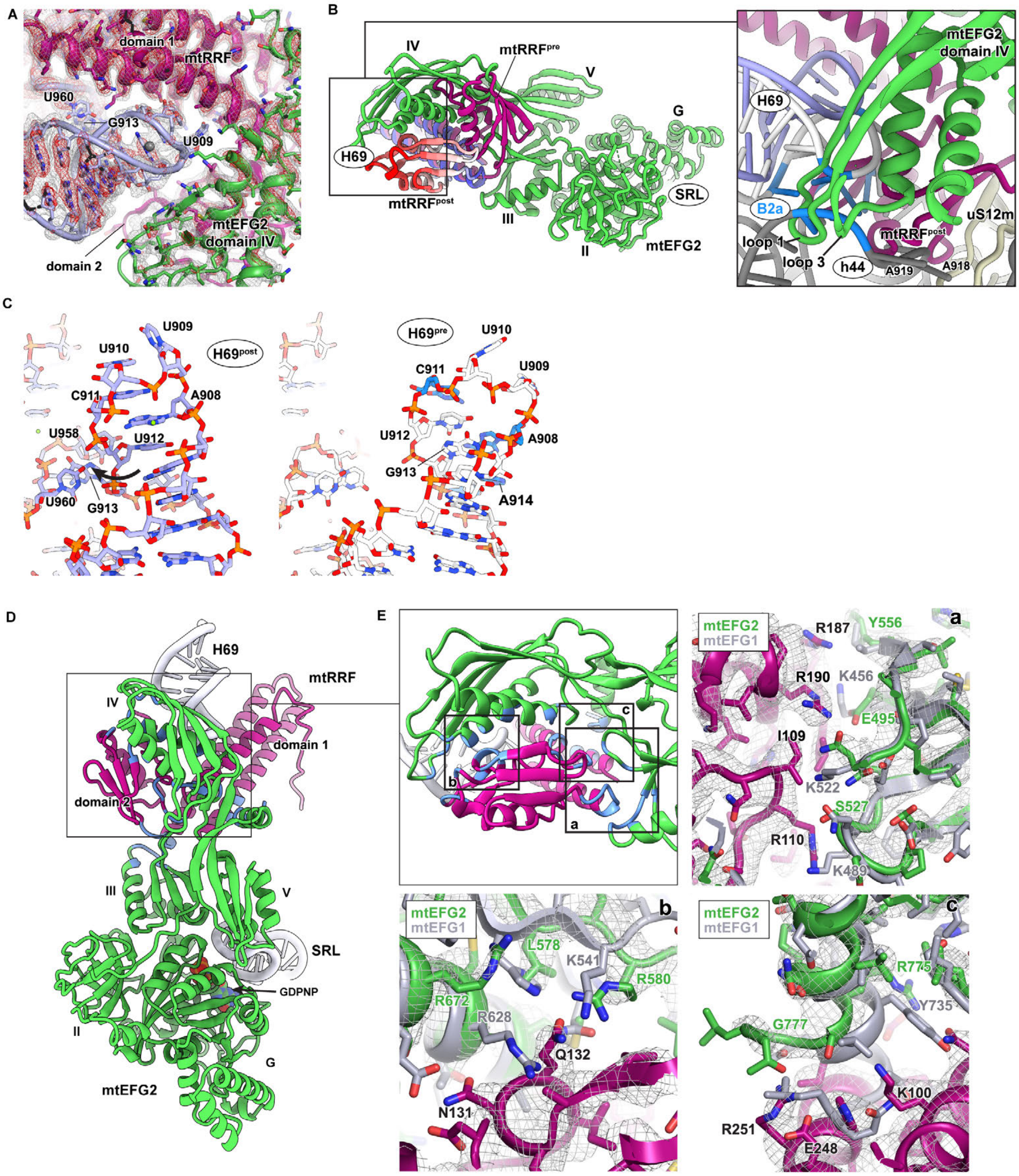


**Figure S4** Interactions of mtEFG2 and mtRRF in the post-splitting complex, Related to Figure 5 and Figure 6

A) 16S rRNA helix 69 (H69) is tightly embedded in the 39S post-splitting complex between domains 1 and 2 of mtRRF (purple). The EM density is shown at two thresholds. G913 of tilted H69 stacks with U958 and U960 of neighboring H71.

B) Motion of mtRRF domain 2 is induced by binding of mtEFG2 (green) to the ribosome in order to avoid clashing with domain IV of mtEFG2. mtRRF of the pre-splitting complex (mtRRF$^{pre}$, purple) and the post-splitting (mtRRF$^{post}$) are shown. mtRRF$^{post}$ has been coloured according to the displacement of the atoms with respect to mtRRF$^{pre}$ as calculated in PyMOL using the draw_rotation_axis.py script (P.G. Calvo). Adjacent ribosomal sites are indicated (SRL = sarcin ricin loop). On the right, ribosomal RNA is shown in white and grey for the LSU and SSU, respectively, of the pre-splitting complex, and in blue for the LSU of the post-splitting complex. Residues contributing to the B2a intersubunit bridge are highlighted in blue. H69 motion disrupts B2a in the post-splitting complex and mtEFG2 domain IV as well as mtRRF domain 2 clash with the SSU 12S rRNA (h44) leading to dissociation of the ribosomal subunits.

C) Comparison of the position and conformation of the nucleotides of 16S rRNA H69 in the 39S post-splitting (light blue) and the 55S pre-splitting (white) complex after superposition of their 16S rRNAs. The rRNA helix is shifted and the stacking of the RNA bases has been altered in the post-splitting complex. Flipping of G913 in between U958 and U960 of H71 is indicated by an arrow. Nucleotides that contribute to the intersubunit bridge B2a are colored in blue in the pre-splitting state.

D) An overview of the complex of mtRRF (purple) and mtEFG2 (green) is shown. Domains of mtEFG2 are indicated in Roman numerals. Molecular interactions between both proteins in the post-splitting complex have been analyzed using the Chimera contact analysis tool at default settings. Amino acids calculated to interact are highlighted (blue). The main interaction area between mtRRF and mtEFG2 domain IV is boxed and a close-up is shown in Figure S4E.

E) The three main interaction sites between mtRRF and mtEFG2 domain IV are indicated by boxes in the first panel. These areas include the interaction of mtEFG2 with the mtRRF linker region as well as of the mtEFG2 C-terminus with mtRRF domain 1. Moreover, close-up views of the boxed areas are depicted with the labeling matching the first panel. Domain IV of mtEFG1 (grey) has been superpositioned onto mtEFG2 (green) to compare the amino acid composition of both proteins in the respective regions. The experimental EM density for mtEFG2 is shown as mesh. In (a) the electrostatic unfavorable amino acid composition of mtEFG1 at the predicted contact site to the mtRRF linker is visible. Moreover, the data indicate a clash between the C-terminal extension of mtEFG1 and mtRRF domain 1 (c).

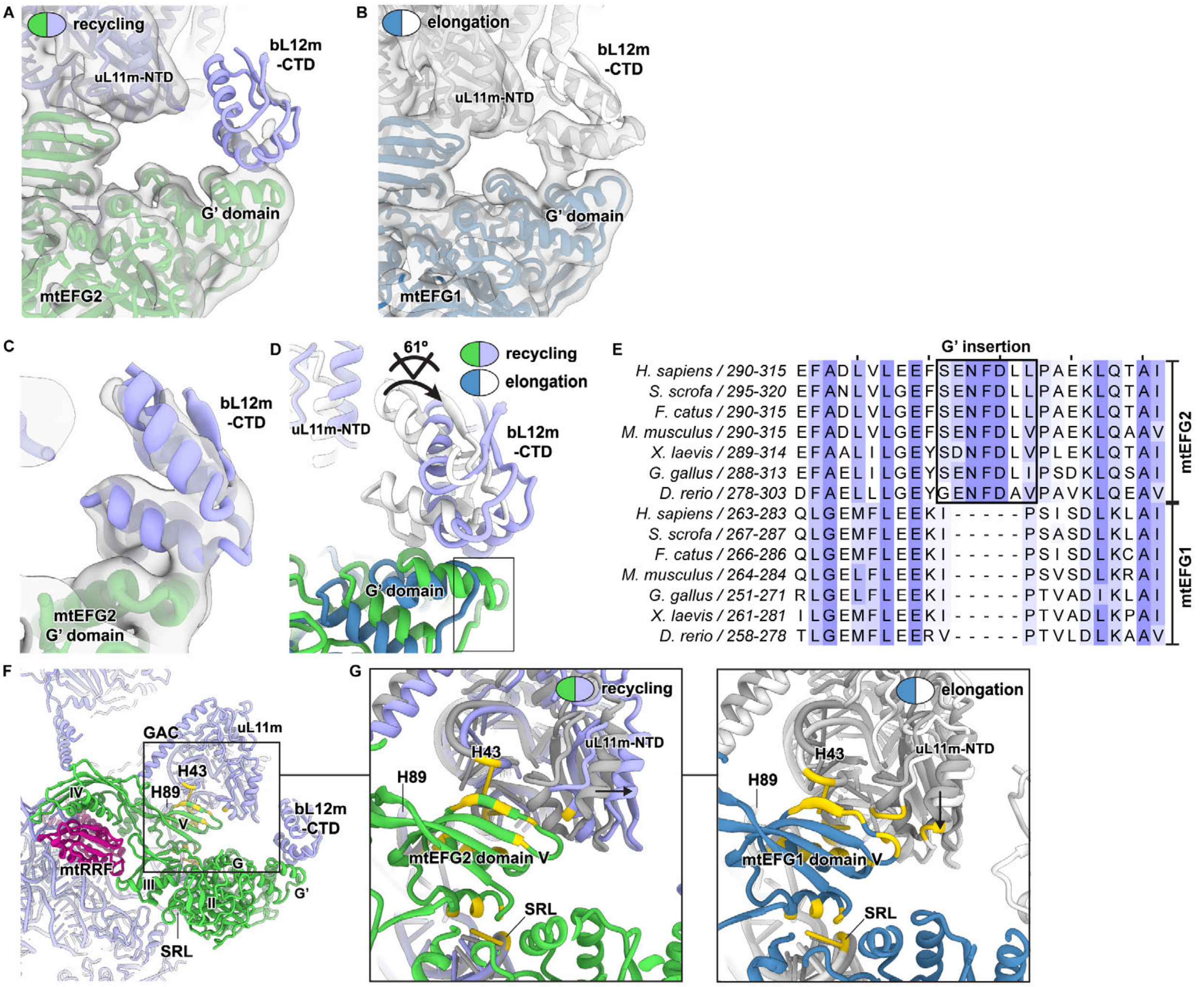


**Figure S5** Interaction of mtEFG2 with the ribosomal GTPase-associated center, Related to Figure 6

A – B) Recruitment of translational GTPases to the ribosome is facilitated by binding of the C-terminal domain of ribosomal bL12m (bL12m-CTD) to the GTPase domains. In mtEFG1 (blue) and mtEFG2 (green), bL12m-CTD interacts with an insertion in the GTPase domain called G'. bL12m-CTD appears to interact more weakly with mtEFG2. EM densities lowpass-filtered to 7 Å for translation elongation (PDB = 6YDP, EMD-10778 (Kummer and Ban, 2020)) and ribosome recycling complexes are shown as semi-transparent surfaces. Thresholds have been adjusted manually such that mtEFG1 and mtEFG2 as well as adjacent ribosomal elements are shown at a similar intensity. Remarkably, at these settings the EM density corresponding to bL12m-CTD is clearly more prominent in the translation elongation complex suggesting a higher occupancy or reduced flexibility of bL12m-CTD on mtEFG1. bL12m-CTD may thus interact more strongly with mtEFG1 than with mtEFG2.

C) Density for the bL12m-CTD in the ribosome recycling complex lowpass-filtered to 7 Å and at a lower threshold than shown in A) permits unambiguous assignment of the orientation of bL12m-CTD on the G' domain of mtEFG2.

D) Superposition of the translation elongation (PDB = 6YDP (Kummer and Ban, 2020)) and ribosome recycling complexes shows that bL12m-CTD adopts different orientations on the G' domains of mtEFG1 (blue, corresponding bL12m-CTD in white) and mtEFG2 (green, corresponding bL12m-CTD in light blue). The degree of rotation has been calculated in PyMOL using the draw_rotation_axis.py script (P.G. Calvo). The different orientation of bL12m-CTD may be caused by an mtEFG2-specific polypeptide insertion (box).

E) Clustal Omega-based sequence alignment of the mtEFG2 insertion with respect to the corresponding region in mtEFG1 has been colored according to conservation in Jalview (overall conservation threshold = 30, conservation threshold for the boxed insertion segment = 50). Species and residue boundaries of the sequences are indicated on the left.

F – G) Interaction of domain V of mtEFG2 (green) and mtEFG1 (blue) with the GTPase-associated center (GAC) that is important for factor binding and GTPase activation. The GAC consists of rRNA helices H43 and H89, uL11m and the sarcin ricin loop (SRL). An overview of the interaction of mtEFG2 with the GAC is given in F) and the box indicates the position of the close-up views shown in G). The GAC moves upon binding of mtEFG2 or mtEFG1, respectively, with regard to its conformation in the factor-free mitoribosome (grey, PDB = 5AJ4 (Greber et al., 2015)), albeit in different directions as visualized by the arrows. Interactions have been calculated in Chimera using default settings and interacting residues are highlighted in yellow. These results show that the number of mtEFG2 interaction with the GAC is lower than for mtEFG1, especially with respect to uL11m.

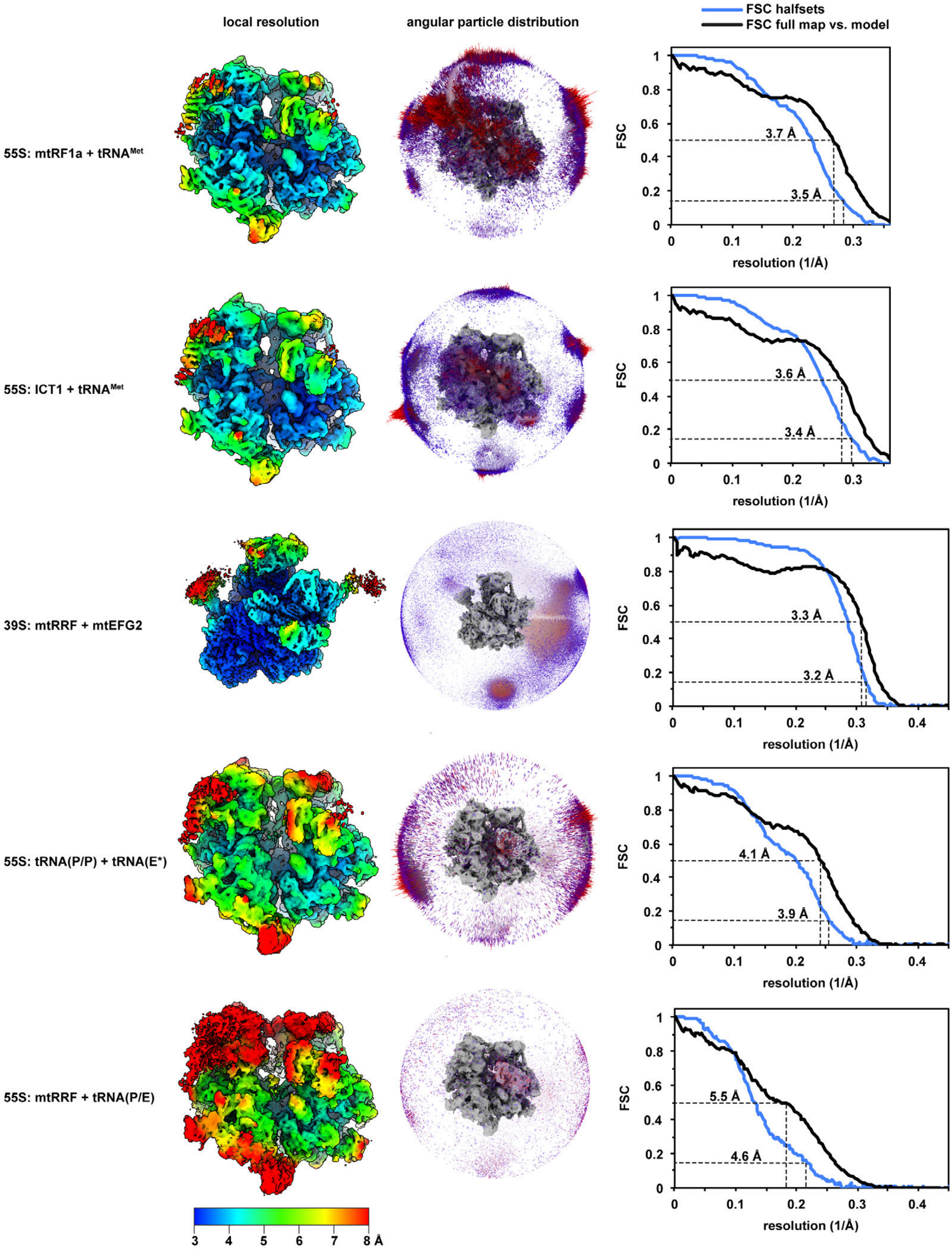


**Figure S6** Local resolution, angular particle distribution and FSC calculation, Related to Figure 4
On the left, the local resolution has been plotted for each complex onto the surface of the Gaussian-filtered EM maps (s.d. corresponds to the pixel size, i.e. 1.39 in case of mtRF1a and ICT1, and 1.087 in case of the

recycling complexes) that have been postprocessed in RELION3.1 before. The local resolution was estimated using the implementation in RELION3.1 and the corresponding color key is provided at the bottom. In the middle, the angular distributions of the particles contained in the final reconstructions are visualized. The ribosome or ribosomal subunit, respectively, is shown in grey in the center for reference. On the right, the FSC curves have been plotted to show the correlations between the data half-sets (blue) and the model vs. the full map (black), respectively. Estimated resolutions at the 0.143 criterion for the half-sets FSC and the 0.5 criterion for the model vs. map FSC are indicated. In case of 55S: mtRRF + tRNA(P/E) the density was too poor for manual model building and the final structural model was derived by rigid body fitting and only mild adjustments of connections and refinement to avoid clashing of the fitted models. The model vs. map FSC deviates in this case somewhat from the resolution estimate of the half-sets. As the rigid bodies for the final model have however been derived from earlier high-resolution structures as detailed in the Methods section, we consider our model valid and attribute this gap to the strong local resolution variability of our map as evident from the local resolution plots.

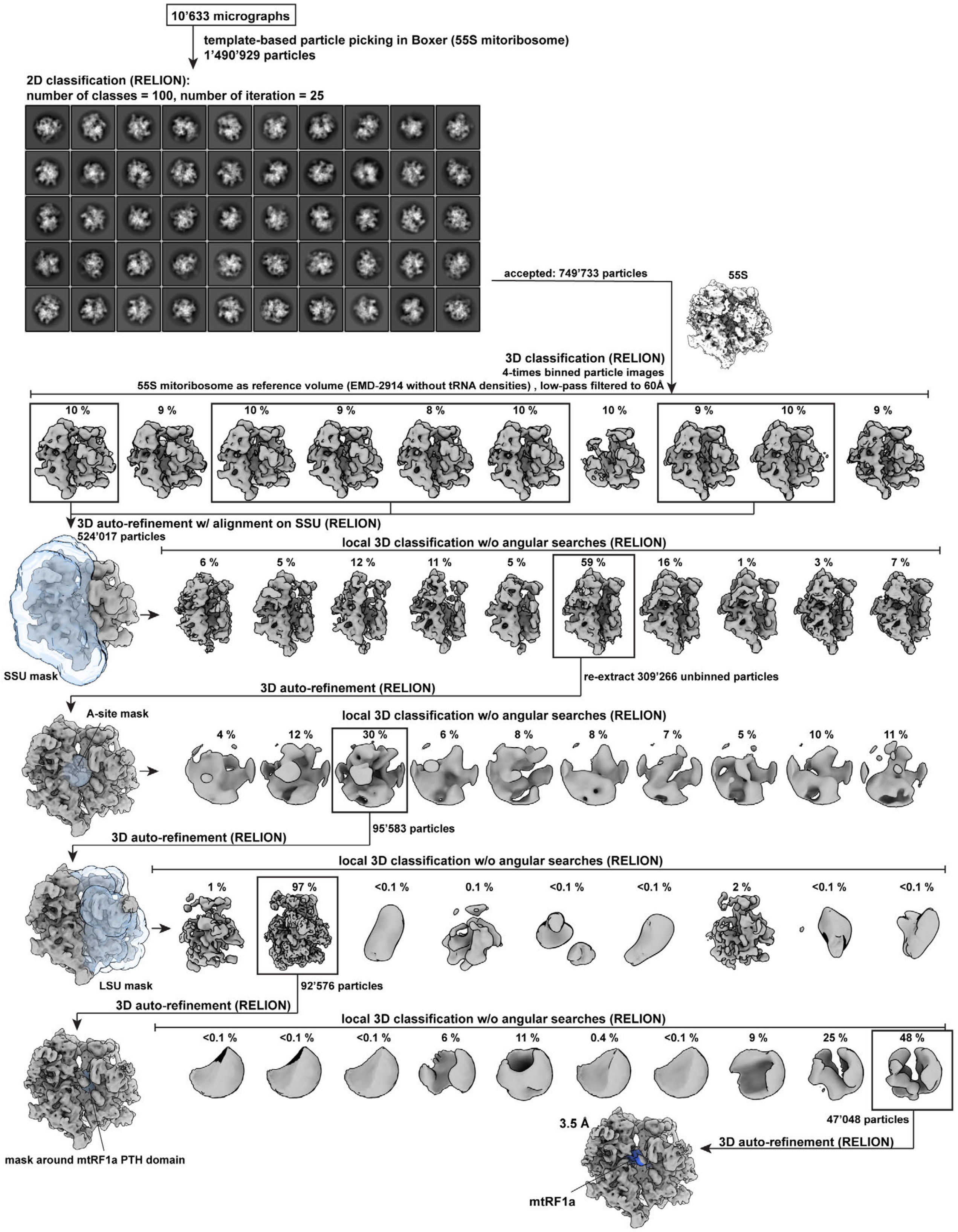


**Figure S7** Classification scheme for the 55S mitoribosome in complex with mtRF1a, Related to Figure 1
The classification scheme is shown that yielded the final 3D reconstruction of the 55S mitoribosome in complex with mtRF1a. On top, only 2D classes are displayed that have been considered for further analysis.

The initial 55S mitoribosome reference model derived from EMD-2914 (Greber et al., 2015) is shown in white. Distribution of particles in distinct classes during 3D classifications is given in %. Masks that have been applied for local 3D classifications and, whereever indicated, for local gold-standard 3D refinements, are shown in blue and semitransparent in the context of the ribosome. mtRF1a is colored in blue in the final reconstruction.

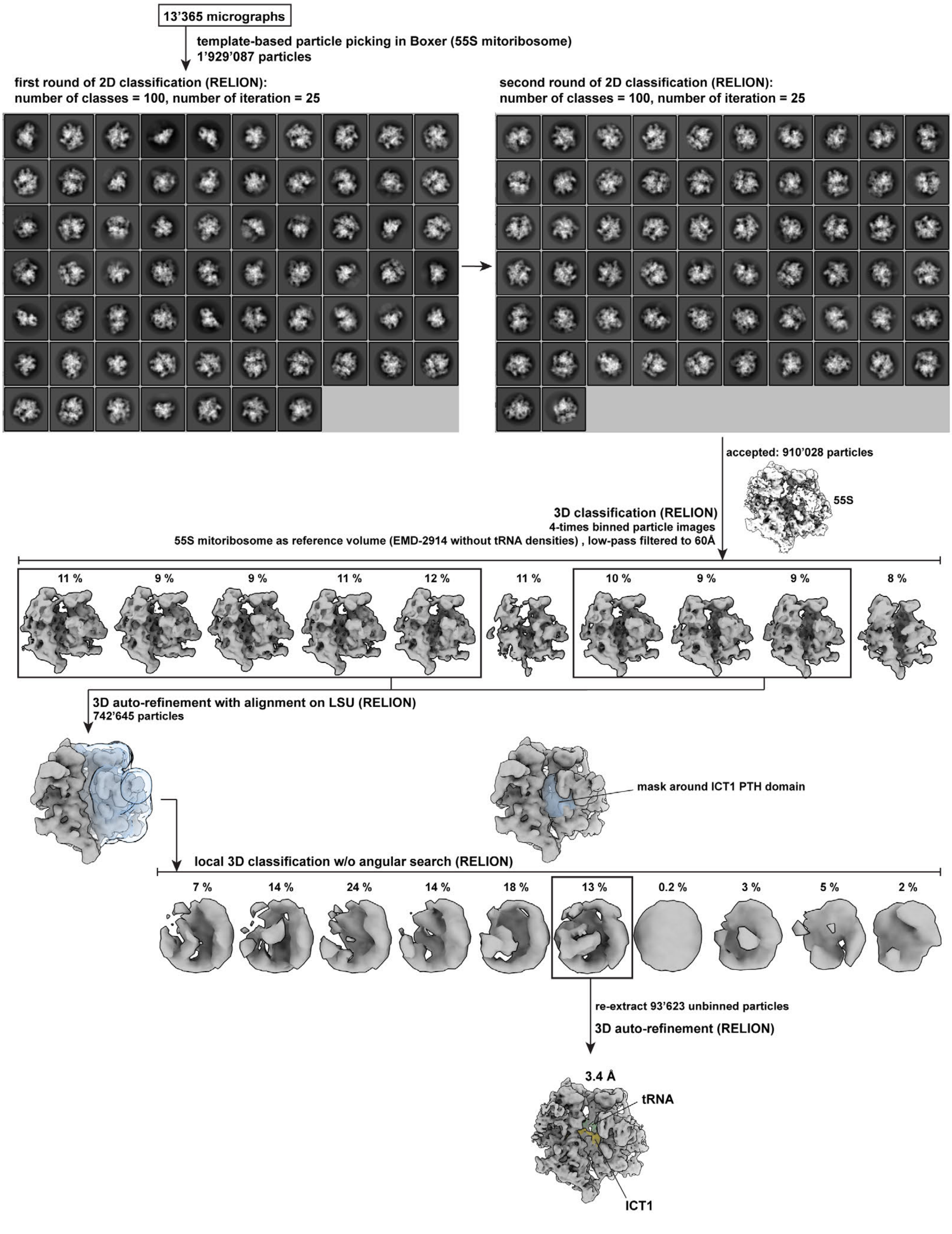


**Figure S8** Classification scheme for the 55S mitoribosome in complex with ICT1, Related to Figure 1

The classification scheme is shown for the final reconstruction of the 55S mitoribosome in complex with ICT1 in a similar fashion than for mtRF1a. On top, only 2D classes are displayed that have been considered for further analysis and the initial 55S mitoribosome reference model derived from EMD-2914 is shown in white. Particle distributions across distinct particle populations that resulted from 3D classifications are given in %. Masks that have been applied for local 3D classifications and, whereever indicated, for local gold-standard 3D refinements are shown in blue and semitransparent in the context of the ribosome. ICT1 is colored in yellow and mtRNA$^{Met}$ in green in the final reconstruction.

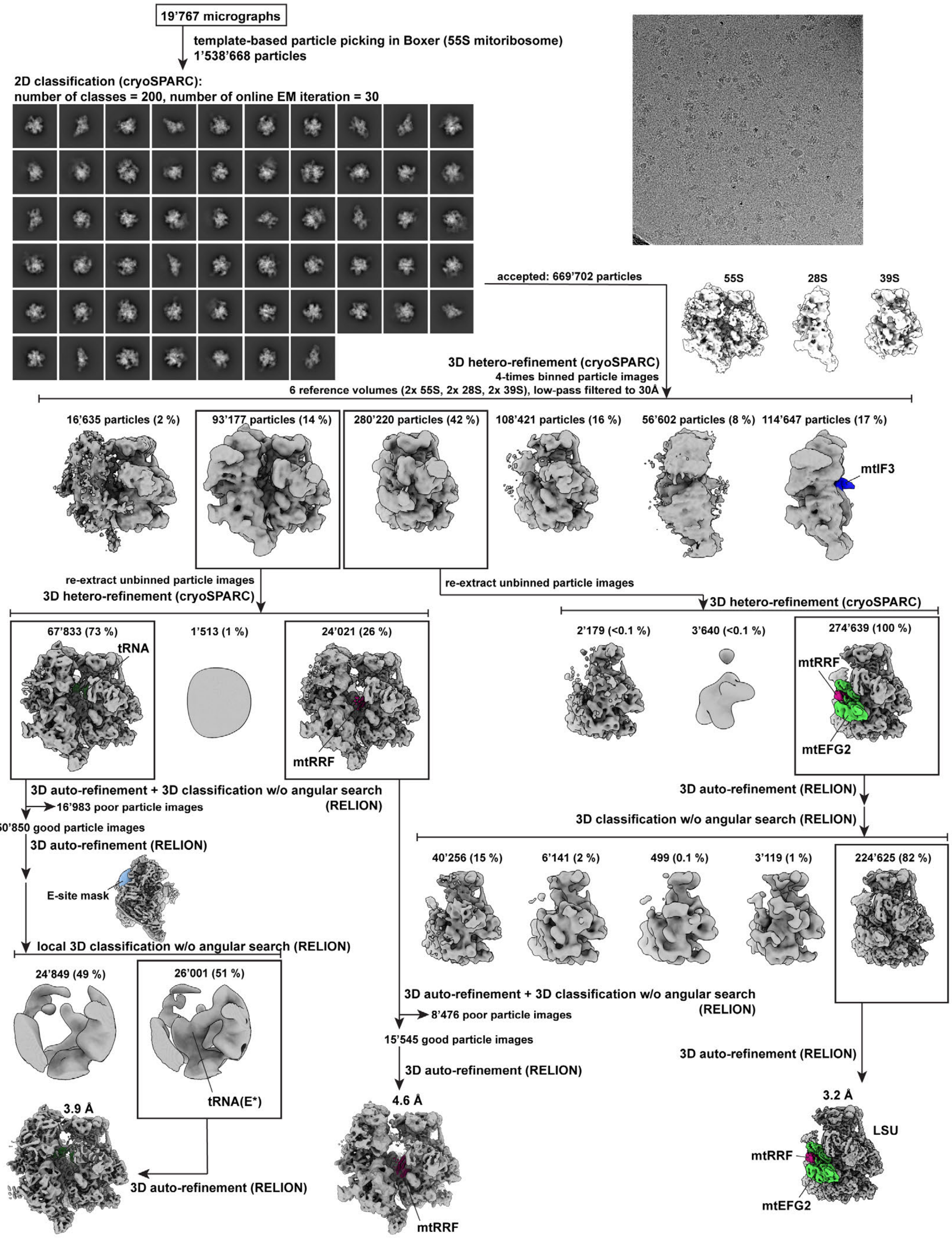


**Figure S9** Classification scheme for the ribosome recycling complexes, Related to Figure 4

The data of the ribosome-recycling sample have been analyzed in cryoSPARC and RELION3.1. It is indicated in the scheme, which step has been carried out in which program. On top, only 2D classes are displayed that have been considered for further analysis and an example micrograph is included. The initial 55S, 39S and 28S mitoribosome reference models derived from EMD-2914 are shown in white. Distributions of the input particles upon 3D heterogeneous refinements in cryoSPARC or 3D classifications in RELION3.1 are indicated as particle number and in percent. The respective factors including mtIF3 (blue), tRNA (dark green), mtRRF (purple) and mtEFG2 (green) in complex with the ribosome have been highlighted in color in the final 3D reconstructions and important intermediate steps.

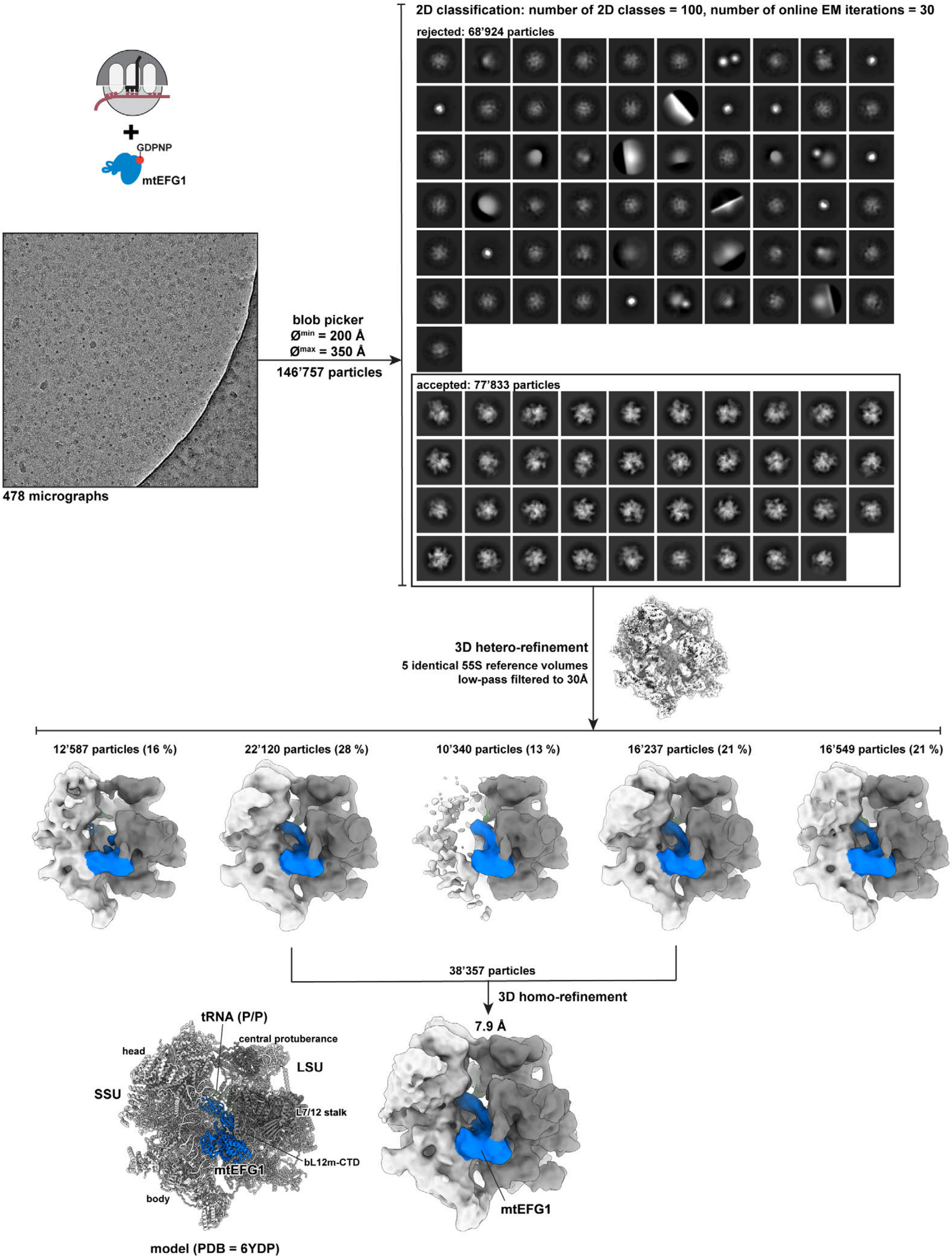


**Figure S10** Classification scheme of the 55S mitoribosome in complex with mtEFG1, Related to Figure 6
The 55S mitoribosome has been incubated with mtEFG1 in the presence of the non-hydrolysable nucleotide analogue GDPNP. Cryo-EM data have been collected on a FEI Tecnai F20 equipped with a Falcon II at 200

kV. An example micrograph is shown. Particles have been picked using the unspecific blob picker in cryoSPARC to avoid any preference in particle selection (minimum particle diameter = 200 Å, maximum particle diameter = 350 Å). Particles were extracted unbinned and subjected to 2D classification in cryoSPARC using 30 online EM iterations and 100 classes. Rejected particle populations as well as accepted ones are shown to visualize that we have selected all particles that reasonably represent the 55S mitoribosome, the 39S LSU, or the 28S SSU. Subsequent heterogeneous refinement in cryoSPARC against the 55S mitoribosome (derived from EMD-2914) as an initial reference volume separates poor 55S particles from intact 55S mitoribosomes. The EM density corresponding to mtEFG1 is already clearly visible in all classes and highlighted in blue. Particles from two classes that corresponded to the best 55S densities were pooled and refined in cryoSPARC to derive the final 3D reconstruction. The corresponding structural model is shown on the left (PDB 6YDP, (Kummer and Ban, 2020).

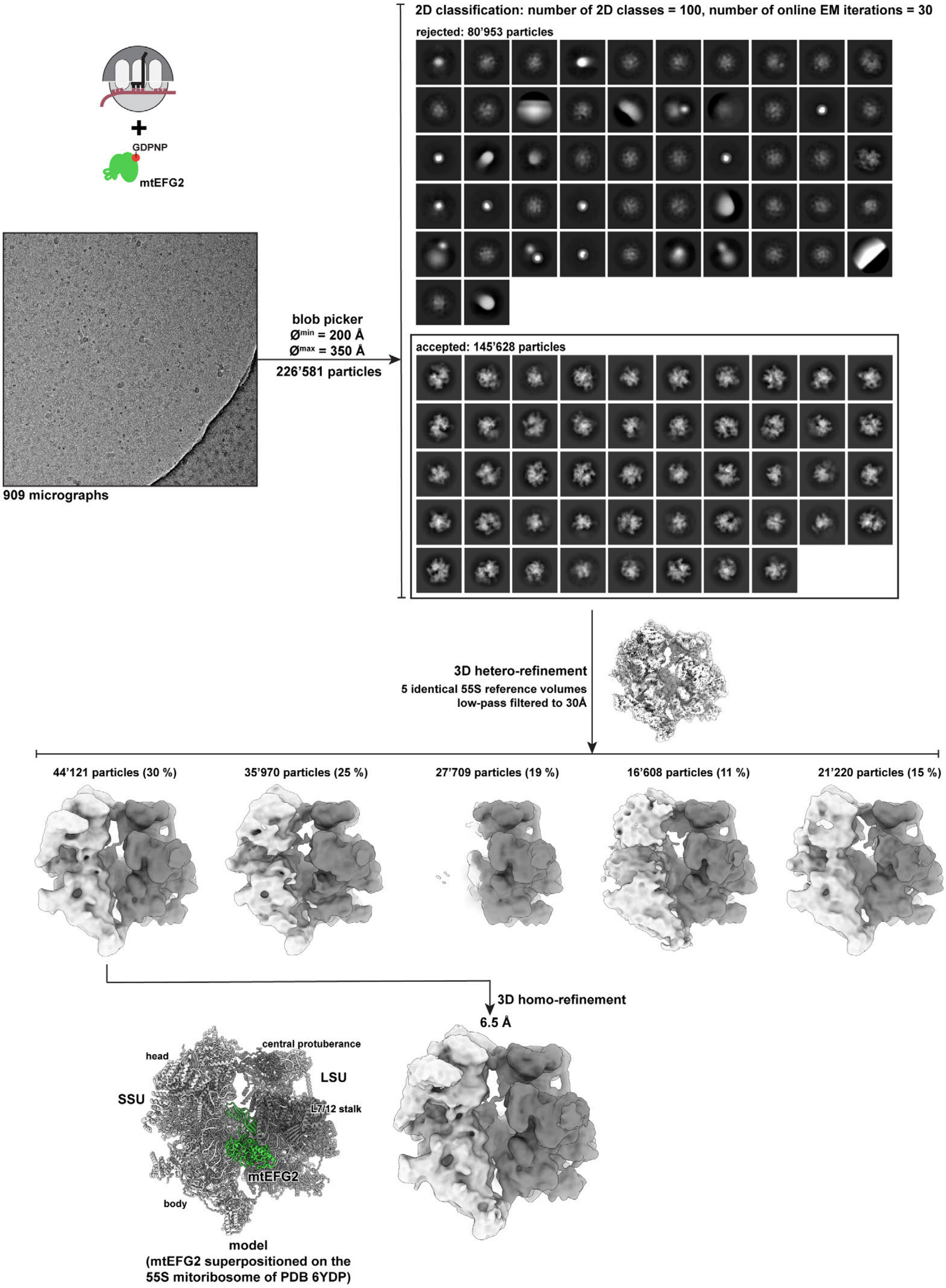


**Figure S11** Classification scheme of the 55S mitoribosome incubated with mtEFG2 and GDPNP, Related to Figure 6

Identical to the mtEFG1 sample shown in Figure S10, mtEFG2 has been incubated with the 55S mitoribosome and the non-hydrolysable nucleotide analogue GDPNP. Cryo-grids were prepared and data were collected on a FEI Tecnai F20 with Falcon II at 200 kV. The data analysis approach was identical to the one described for mtEFG1 in Figure S10 to allow direct comparison of the two datasets. In brief, particles were picked in an unbiased fashion using the blob picker in cryoSPARC. All class averages derived from subsequent 2D classification (30 EM iterations, 100 classes) are shown. Particle populations are indicated that have been retained for further 3D heterogeneous refinement against the 55S mitoribosome as initial reference. None of the classes contains density for mtEFG2. The population representing the best-resolved 55S mitoribosome has been used for homogeneous refinement in cryoSPARC to derive the final 3D reconstruction. On the left, the 55S mitoribosome is shown as a structural model and mtEFG2 has been superpositioned on it by aligning the 39S: mtRRF + mtEFG2 post-splitting complex with the 16S rRNA of the 55S mitoribosome. In this model, mtEFG2 is highlighted in green to indicate its potential position on the 55S mitoribosome. However, corresponding density is missing in the experimental 3D reconstruction showing that mtEFG2 cannot bind the mitoribosome in the absence of mtRRF, in contrast to mtEFG1.

### Table S1. EM data collection and refinement statistics

| | | | | | |
|---|---|---|---|---|---|
| **EM data collection:** All datasets have been collected on a FEI Titan Krios with Falcon 3 DED. | | | | | |
| Number of micrographs | 10’633 | 13’365 | 19’767 | | |
| Pixel size (Å) | 1.39 | 1.39 | 1.087 | | |
| Defocus range (µm) | 0.5 – 2 | 1 – 2.5 | 1 – 2.7 | | |
| Voltage (kV) | 300 | 300 | 300 | | |
| Electron dose (e-/Å$^2$) | 40 | 40 | 40 | | |
| *Name of 3D reconstruction* | 55S: mtRF1a + tRNA$^{Met}$ | 55S: ICT1 + tRNA$^{Met}$ | 39S: mtRRF + mtEFG2 | 55S: mtRRF + tRNA(P/E) | 55S: tRNA(P/P) + tRNA(E*) |
| EMDB map entry | EMD-12527 | EMD-12529 | EMD-12567 | EMD-12568 | EMD-12569 |
| PDB coordinate entry | PDB 7NQH | PDB 7NQL | PDB 7NSH | PDB 7NSI | PDB 7NSJ |
| Final number of particles | 47’048 | 93’623 | 224’731 | 15’545 | 26’001 |
| Resolution (Å) (FSC = 0.143) | 3.5 | 3.4 | 3.2 | 4.6 | 3.9 |
| Map sharpening B factor | -100.86 | -98.1415 | 43.5792 | 17.0027 | 17.9615 |
| **Refinement and validation statistics:** Real-space refinement in PHENIX | | | | | |
| Overall model geometry | | | | | |
| clashscore (all atoms) | 4.18 | 9.48 | 7.72 | 8.16 | 7.90 |
| MolProbity score | 1.43 | 1.70 | 1.71 | 1.71 | 1.67 |
| rmsd (bonds) | 0.004 | 0.002 | 0.002 | 0.002 | 0.002 |
| rmsd (angles) | 0.916 | 0.543 | 0.515 | 0.477 | 0.467 |
| Ramachandran plot (%) | | | | | |
| favoured | 96.53 | 96.82 | 95.83 | 97.13 | 97.13 |
| allowed | 3.41 | 3.18 | 4.15 | 2.85 | 2.86 |
| outliers | 0.06 | 0.01 | 0.02 | 0.02 | 0.01 |
| Rotamer outliers (%) | 0.06 | 0.03 | 0.02 | 1.37 | 1.30 |
| Cβ outliers (%) | 0.03 | 0.00 | 0.00 | 0.00 | 0.00 |
| Peptide plane (%) | | | | | |
| cis proline / general | 0.8/0.0 | 0.8/0.0 | 1.1/0.0 | 0.8/0.0 | 0.8/0.0 |
| twisted proline / general | 0.1/0.0 | 0.1/0.0 | 0.0/0.0 | 0.1/0.0 | 0.0/0.0 |
| CaBLAM outliers (%) | 1.71 | 1.55 | 1.90 | 1.55 | 1.57 |
| Validation RNA | | | | | |
| Sugar pucker outliers (outlier/total) | 6/2651 | 3/2648 | 1/1611 | 5/2647 | 10/2719 |
| CC (mask) | 0.8 | 0.77 | 0.82 | 0.74 | 0.83 |
| CC (box) | 0.75 | 0.62 | 0.63 | 0.71 | 0.78 |
| Resolution estimates | | | | | |
| FSC (half maps; 0.143) | 3.5 | 3.4 | 3.2 | 4.6 | 3.9 |
| FSC (model vs. full map; 0.5) | 3.7 | 3.6 | 3.3 | 5.5 | 4.1 |
